# Machine-learning from *Pseudomonas putida* Transcriptomes Reveals Its Transcriptional Regulatory Network

**DOI:** 10.1101/2022.01.11.475908

**Authors:** Hyun Gyu Lim, Kevin Rychel, Anand V. Sastry, Joshua Mueller, Wei Niu, Adam M. Feist, Bernhard O. Palsson

## Abstract

Bacterial gene expression is orchestrated by numerous transcription factors (TFs). Elucidating how gene expression is regulated is fundamental to understanding bacterial physiology and engineering it for practical use. In this study, a machine-learning approach was applied to uncover the genome-scale transcriptional regulatory network (TRN) in *Pseudomonas putida*, an important organism for bioproduction. We performed independent component analysis of a compendium of 321 high-quality gene expression profiles, which were previously published or newly generated in this study. We identified 84 groups of independently modulated genes (iModulons) that explain 75.7% of the total variance in the compendium. With these iModulons, we (i) expand our understanding of the regulatory functions of 39 iModulon associated TFs (e.g., HexR, Zur) by systematic comparison with 1,993 previously reported TF-gene interactions; (ii) outline transcriptional changes after the transition from the exponential growth to stationary phases; (iii) capture group of genes required for utilizing diverse carbon sources and increased stationary response with slower growth rates; (iv) unveil multiple evolutionary strategies of transcriptome reallocation to achieve fast growth rates; and (v) define an osmotic stimulon, which includes the Type VI secretion system, as coordination of multiple iModulon activity changes. Taken together, this study provides the first quantitative genome-scale TRN for *P. putida* and a basis for a comprehensive understanding of its complex transcriptome changes in a variety of physiological states.

## 1. Introduction

*Pseudomonas putida* has received much attention as a workhorse for numerous biotechnological applications due to its versatile metabolism that allows it to utilize a broad spectrum of substrates and express stress-tolerant phenotypes^1,2^. Despite its industrial importance, understanding of its genome-scale transcriptional regulatory network (TRN) is limited compared to other widely used microorganisms in biotechnology, such as *Escherichia coli* and *Bacillus subtilis*. Therefore, unveiling *P. putida*’s TRN is imperative to improving our mechanistic understanding of gene expression changes to maximize its biotechnological potential^3,4^.

Traditionally, genome-scale TRNs have been reconstructed in a bottom-up fashion by revealing regulons, one at a time, which are defined as the set genes regulated by the same transcription factor (TF)^5^. Specifically, the binding of TFs to putative promoter sites was historically investigated *in vitro* using low-throughput experimental techniques such as an electrophoretic mobility shift assay^6^ and a fluorescence polarization assay^7,8^. Recently, with the development of new sequencing technologies that use chromatin immunoprecipitation (ChIP), genome-wide *in vivo* binding studies became possible^9,10^. These results are typically compared with differentially expressed genes from TF knock-out and overexpression studies to distinguish functional binding sites from nonfunctional ones. Although these methods are essential to identify and validate regulons, elucidation of the genome-scale TRN, focusing on one known TF at a time is very laborious as there are approximately 500 proteins involved in transcriptional regulation in *P. putida* (Supplementary Data). Thus, a broader strategy is needed to elucidate the global TRN.

Recently, independent component analysis (ICA) has been shown to efficiently uncover bacterial genome-scale TRNs from gene expression profiles, obtained by using methods such as microarray hybridization and RNA sequencing^11–13^. ICA is an unsupervised blind source separation algorithm, which can identify statistically independent components (ICs) from mixed signals^14–16;^ it was found to perform the best in a comparative analysis of signal extraction algorithms for TRN inference^17^. ICs from gene expression profiles are independently modulated groups of genes (iModulons), which are likely under the regulation of the same underlying biological signal (e.g. TF or genetic perturbation). iModulons are similar to regulons^18^, but not identical since they were obtained in a top-down data-driven manner. While Principal Component Analysis (PCA) is often used to identify variance within datasets, principal components (PCs) are statistical measures that usually lack clear biological interpretations. However, we have demonstrated that iModulons recapitulate known regulatory mechanisms and can thus be considered as knowledge-based representations of the composition of the transcriptome. Indeed, recent studies successfully applied ICA for model bacteria (e.g., *E. coli*^13^, *Bacillus subtilis*^11^) and even less-studied bacteria (e.g., *Staphylococcus aureus*^12^, *Mycobacterium tuberculosis*^19^*, Sulfolobus acidocaldarius*^19^, *Pseudomonas aeruginosa*^20^) to unveil their genome-scale TRNs.

In this study, to unveil the genome-scale TRN of *P. putida*, we perform ICA for an RNA-seq compendium, named *putida*PRECISE321, consisting of 321 high-quality gene expression profiles, of which 305 were previously published and 16 were newly generated in this study (**Supplementary Data**). We obtained 84 iModulons, which explain 75.7% of the variation in the compendium. With the iModulons, we show that the genome-scale TRN in *P. putida* can be effectively uncovered by identifying regulatory target genes of 38 TFs. In cases where the iModulon did not match previously described regulons well, we propose improved gene-regulator interactions that are corroborated by motif analysis. In addition, we demonstrate the ability of iModulons to comprehensively interpret transcriptional responses to environmental or evolutionary changes. We demonstrate the iModulon clustering, effectively representing a stimulon, to suggest an activating condition of a less-characterized functional cluster (i.e., the Type VI Secretion System). Collectively, the 84 iModulons delineate the genome-scale TRN in *P. putida* by deconvolution of complex transcriptomic responses orchestrated by multiple TFs. Detailed, searchable, interactive dashboards for all 84 iModulons are publicly available on iModulonDB (https://imodulondb.org)^21^.

## 2. Results

### 2.1. ICA reveals 84 independently modulated groups of genes in *P. putida*

We generated a compendium of RNA-seq datasets for *P. putida* by downloading, aligning, and performing rigorous quality control on all public datasets available from SRA using our established pipeline^18^ (See **Methods**; **Fig. 1a** and **Supplementary Fig. 1**). We named the compendium *putida*PRECISE321, for *putida* Precision RNA-seq Expression Compendium for Independent Signal Exploration. It consists of 321 samples from 118 unique experimental conditions across 21 projects^22–31^. We centered each project to a baseline condition in order to remove batch effects (**Supplementary Fig. 2**) and then used the ICA algorithm^18^ to extract all independent expression signals (iModulons) from the data. The results, therefore, represent co-regulated gene sets and regulator activities across all genes perturbed by any available high-quality transcriptome for *P. putida*.

**Figure 1.**
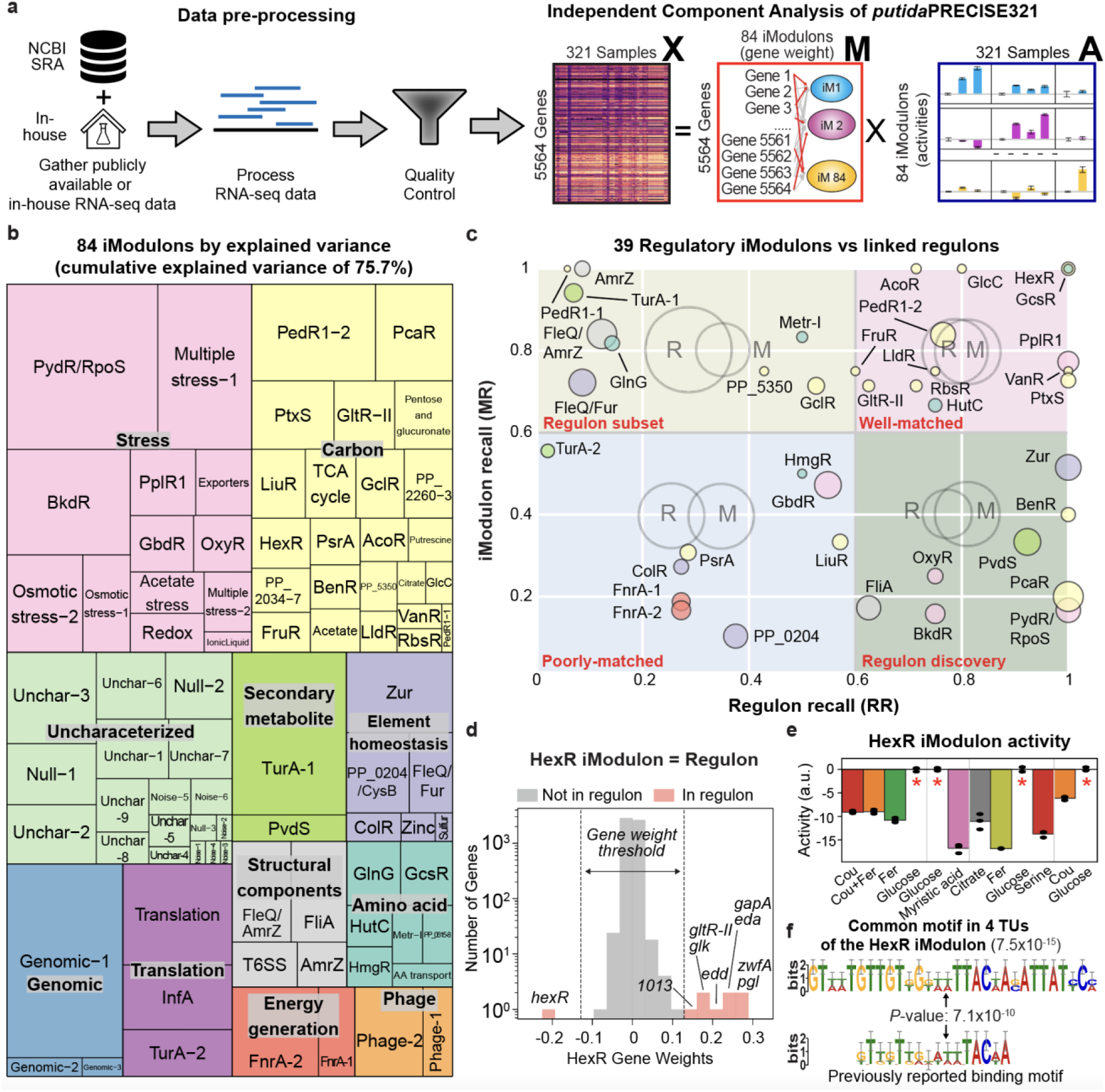
Independent component analysis (ICA) of the *putida*PRECISE321 transcriptome compendium. (**a**) Overall data processing pipeline for independent component analysis (ICA) of *putida*PRECISE321. By performing ICA, the gene expression profiles [**X**] (5564 genes by 321 conditions) were deconvoluted into an [**M**] matrix containing weights of 5564 genes for each of the 84 iModulon identified, and an [**A**] matrix containing iModulon activities for each condition. A part of this diagram was adapted from Anand et al., 2021^18^. (**b**) A treemap of *P. putida*’s 84 iModulons. The size of each rectangle indicates the explained variances in the data sets attributable to each iModulon. (**c**) Bottom-up vs top-down assessment of co-regulated genes (i.e, bio-molecular vs data-driven). The scatter plot shows Regulon Recalls (RR) and iModulon Recalls (IR) for 39 regulatory iModulons. For iModulons with names consisting of two regulators, RR and IR are the values of the first regulator. Colors indicate iModulon types. (**d**) An example of a well-matched case: HexR. Gene weights of the HexR iModulon. (**e**) Condition-dependent HexR iModulon activities with the supplementation of different carbon sources. Asterisks indicate the reference conditions (glucose minimal media). It should be noted that each project (separated by gray lines) was normalized to its unique control condition (**Supplementary Data**) to remove batch effects, so activity values from different projects should not be directly compared. (**f**) Enriched motif (top) from the upstream of HexR-regulating genes and previously reported HexR binding site (bottom, obtained from CollectTF^65^).

The ICA of *putida*PRECISE321 revealed 84 iModulons. Each iModulon has member gene weights (stored in the columns of a matrix **M**), from which outliers are identified as “member genes”, and activities in each sample, or condition (stored in the rows of a matrix **A**). The 84 iModulons contain genes with closely related biological functions (**Fig. 1b** and **Supplementary Fig. 3a**). The median gene number in the 84 iModulons was 10 (**Supplementary Fig. 3b**). Together, these iModulons include 1265 genes (23% of the total annotated ORFs), whose expression levels actively changed under the conditions represented in the dataset. A total of 75.7% of the gene expression variance was mechanistically explained by these iModulons, which is comparable to the variance found by PCs (**Supplementary Fig. 3c**).

The iModulons were characterized and categorized into four groups, depending on their level of characterization and likely underlying mechanisms: regulatory, functional, genomic, and uncharacterized iModulons. “Regulatory” iModulons (46.4% of the 84 iModulon) have significant overlaps (false discovery rates, FDRs < 10^−4^) with previously reported regulons mostly from bottom-up experiments. We compiled 1,993 experimentally validated or predicted interactions, as part of this study, from multiple databases and literature (see **Methods** and **Supplementary Data**). Consequently, we found 39 iModulons associated with TFs (**Supplementary Table 1**). In addition, 24 “functional” iModulons (28.6%) were defined by enrichment of genes found in a given metabolic pathway from the KEGG database, or by inferring roles based on gene functional annotations or their active conditions. Additionally, three “genomic” iModulons (3.5%) contain differentially expressed genes due to genomic variations (e.g., gene deletion) in the host strains. The remaining 18 iModulons (21.4%) were classified as “uncharacterized”. They typically contained short hypothetical genes or genes with unknown functions or unclear active expression conditions. “Uncharacterized” iModulons represent potential discoveries of new regulons. Regulatory or functional iModulons were named based on their regulators (if associated) or related characteristics (see **Methods**). Genomic or uncharacterized iModulons were numbered in order based on the number of genes that they contain, with smaller iModulons appearing first.

To investigate the accuracy of TRN uncovered by iModulons, we systematically compared the 39 regulatory iModulons with associated TF regulons using two metrics (**Fig. 1c**): iModulon recall (MR) and regulon recall (RR), where MR is the fraction of shared genes in a given iModulon and RR is the fraction of shared genes in a linked TF regulon (**Supplementary Fig. 4a**). Depending on these two metrics, regulatory iModulons were further categorized into four groups (one for each quadrant of **Fig. 1c**) with a threshold value of 0.6 for both MR and RR: well-matched, regulon subset, regulon discovery, and poorly-matched^13,32^. 12 regulatory iModulons (both MR and RR ≥ 0.6), including HexR, GlcC, AcoR, and BenR, were “well-matched” with linked regulons (**Supplementary Fig. 4b**), indicating that ICA successfully recapitulated previously reported regulation. Regulon subset iModulons (MR ≥ 0.6 and RR < 0.6) indicate that a subgroup of a linked regulon shows an independent signal, suggesting different strength of regulation within regulon genes or another layer of regulation by unstudied co-regulator(s). Larger established regulons often contain multiple iModulons^13,32^, underlining the fact that regulons have a ‘static’ definition (i.e., TF binding) and iModulons have a ‘dynamic’ definition (i.e., are computed from expression profiles).

We identified ten regulon subset iModulons associated with TFs, including FleQ (PP_4373), GlnG (PP_5048), and PedR1 (PP_2665), known to regulate relatively large numbers of genes. In addition, we also obtained eight “regulon discovery” iModulons (MR < 0.6 and RR ≥ 0.6). These iModulons include most reported target genes of linked TFs but also contain additional genes whose regulation had not been previously reported. Given the similar expression pattern of genes in the same iModulons, these additional genes are potentially new target genes of a linked TF. Alternatively, entire iModulon genes can be co-regulated by a less-studied or yet unidentified regulator, closely associated with a linked TF. Lastly, we obtained eight “poorly-matched” iModulons (both MR and RR < 0.6) which have small overlaps with linked closest regulons. The relatively low regulon overlap of these iModulons may be explained by additional, unknown TFs, weak or variable binding strengths for TFs with large regulons, or incompletely described regulons.

The 84 iModulons reproduce many known details of the *P. putida* TRN, including the TFs and their target genes, regulatory modes, and binding sites. For example, the HexR iModulon has an MR and RR of 1, indicating that it precisely captured the previously reported regulation of HexR (PP_1021), a glucose metabolism regulator^33^ for nine target genes (*zwfA*, *pgl*, *gapA*, *eda*, *edd*, *gltR-II*, *glk*, PP_1013, *hexR*, **Fig. 1d**). Since HexR represses the expression of itself and the 8 other genes^34^, ICA assigned a negative gene weight to *hexR* and a positive gene weight to the remaining genes. In addition, the activity of the HexR iModulon was found to be decreased when non-glucose carbon sources were provided (**Fig. 1e**), consistent with its expected function. Lastly, a binding site of HexR can be also computationally predicted by enriching a consensus motif in the upstreams of iModulon member genes^35^. With a maximum window of 30 bp, we found the “GTWWTGTTGTKGKWWTTACWASATTATYCM” motif is present in the upstream regions of all four transcriptional units (TUs, *e*-value of 7.5×10^−15^, **Fig. 1f,** see **Supplementary Note 2** for each symbol), which perfectly includes a previously reported motif. Similar characterization can be extended to other TFs from iModulons; we provided summaries and identified motifs of selected iModulons in **Supplementary Tables 1 and 2**, respectively. The agreement between well-matched regulons and iModulons shows commonality in the output of the complementary bottom-up (biomolecular) and top-down (data analytics) approaches.

### 2.2. iModulons allows data-driven curation of the TRN in *P. putida*

Regardless of iModulon types, differences between iModulons and corresponding regulons indicate the presence of gaps between the ICA-defined gene groups and existing knowledge and thus allow further data-driven curation of *P. putida*’s TRN. As examples, we provided detailed characterizations of FleQ/Fur, Zur, PP_0204/CysB iModulons, a representative iModulon of each ‘regulon subset,’ ‘regulon discovery,’ and ‘poorly-matched’ iModulons.

*A regulon subset iModulon shows co-regulation in the FleQ regulon by multiple TFs* - ICA identified a regulon subset iModulon (originally named as FleQ-1 but renamed to FleQ/Fur) in the FleQ regulon (MR and RR of 0.72 and 0.09, **Fig. 2a**). FleQ (PP_4373) is a c-di-GMP(cyclic-di-GMP)-dependent TF, known to regulate up to 300 genes, involved in iron transport, flagellar synthesis, and biofilm formation^36,37^. This iModulon contains 36 genes, mostly involved in inorganic ion transport and metabolism (**Supplementary Fig. 5a** and **5b**) and regulated by FleQ. Its activity increased when *P. putida* was grown in a relatively less-aerated condition, in a medium containing acetate, or in cells which were adapted to minimal media conditions; conditions which likely affected oxygen or iron availability (**Fig. 2b**). We hypothesized that this independent signal within the FleQ regulon is owing to the presence of another regulator(s) for genes.

**Figure 2.**
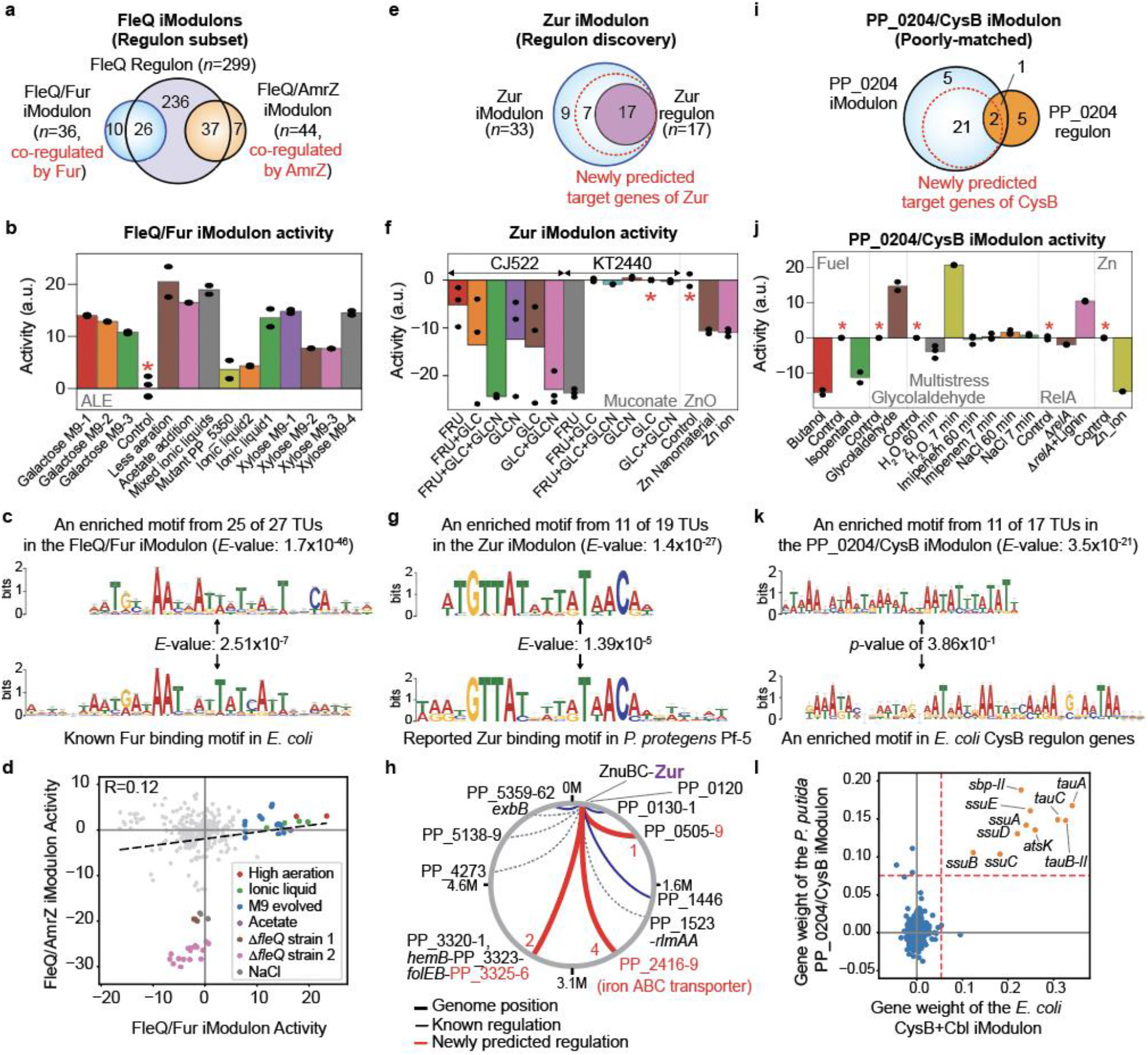
iModulons allow data-driven curation of the TRN in *P. putida*. (**a-d**) Characteristics of the FleQ regulon subset iModulons. (**a**) A Venn diagram of the two FleQ iModulons, FleQ/Fur and FleQ/AmrZ, and the FleQ regulon. (**b**) An activity plot of the FleQ/Fur iModulon in selected conditions. (**c**) Comparison of a motif found in the upstream regions of the FleQ/Fur iModulon genes and previously known binding motifs of Fur in *E. coli* (obtained from CollectTF^65^). The common motif was enriched from 25 out of 27 TUs (94.4% of the genes) in the FleQ/Fur iModulon. (**d**) A scatter plot of activities of the FleQ/Fur and FleQ/AmrZ iModulons for all conditions. The Pearson correlation of the two iModulon activities was 0.12. Selected samples with high activities were colored and growth conditions were indicated in the inset. Activities of the FleQ/AmrZ iModulon changed when *fleQ* was deleted or an excess amount of NaCl was added. (**e-h**) Characteristics of the Zur regulon discovery iModulon. (**e**) A Venn diagram of the Zur iModulon and related Zur regulon. (**f**) An activity plot of the Zur iModulon in selected conditions. High activities of the Zur iModulon were observed when zinc ion or nanomaterial was added. The CJ522 strain is an engineered *P. putida* KT2440 strain Δ*catRBCA*::Ptac:*catA* Δ*pcaHG*::Ptac:*aroY*:*ecdB*:*asbF* Δ*pykA*::*aroG*^D146N^:*aroY*:*ecdB*:*asbF* Δ*pykF* Δ*ppc* Δ*pgi*-1 Δ*pgi*-2 Δ*gcd*^66^. (**g**) Comparison with an enriched motif from 11 TUs in the Zur iModulon (66.7% of the total genes) and a previously reported *P. protegens* Pf-5 Zur binding motif (obtained from CollectTF^65^). (**h**) A schematic diagram of regulation by Zur. Previously known regulations were colored in black while newly predicted regulations were colored in red. Potential indirect regulations were indicated with dashed gray lines. (**i**-**l**) Characteristics of the PP_0204 poorly-matched iModulon. (**i**) A Venn diagram of the PP_0204/CysB iModulon and PP_0204 regulon. (**j**) An activity plot of the PP_0204 iModulon in selected conditions. Asterisks indicate the reference conditions. (**k**) Comparison with an enriched motif from 11 TUs in the PP_0204 iModulon and an enriched motif from 14 TUs in the *E. coli* CysB regulon (obtained from RegulonDB^5^) (**l**) Gene weight correlation of the PP_0204/CysB iModulon and the Cbl+CysB iModulon of *E. coli.* The dashed lines indicate the threshold values (0.054 and 0.075) for *E. coli* and *P. putida* to determine iModulon member genes. See Methods for the calculation. (**a, e,** and **i**) Gene numbers were shown in these diagrams. (**b, f,** and **j**) Asterisks indicate the reference conditions.

To test this hypothesis, we computationally enriched a consensus sequence from the upstream sequences of 27 transcriptional units in this iModulon using MEME^35^. Notably, one motif (AATGYKAAYRATWHTBAYTHBCAWTWD, **Fig. 2c**) was enriched with high confidence (an *e*-value of 0.7×10^−46^) from 25 TUs (34 of the total 36 genes, 94.4%, shown in bold in **Supplementary Fig. 5a**). By comparing this enriched motif to known TF binding sites in public databases using TOMTOM^38^ (See Methods), we found that the motif is highly similar to the binding site of *E. coli* Fur (a ferric uptake regulator) at an *e*-value of 2.51×10^−7^. This result strongly suggests the co-regulation by Fur (PP_4730) in addition to FleQ. Additionally, we obtained another FleQ regulon subset iModulon (MR of 0.86 and RR of 0.12, originally named as FleQ-2 but renamed to FleQ/AmrZ) containing 44 genes, mostly encoding machinery for cell motility such as flagella synthesis (**Fig. 2a** and **Supplementary Fig. 5c**). Its activity decreased when *fleQ* was deleted or cells were osmotically stressed (**Fig. 2d**). Despite their common regulation by FleQ, activities of the two iModulons did not have a significant correlation (Pearson coefficient < 0.12), suggesting the presence of another co-regulator. Indeed, from the upstream regions of 13 out of the 18 TUs (30 of the total 44 genes, 68.2%), another motif (GGTDTTGGYSWTGGCVTYGGCCWGS), similar to the binding site of *P. aeruginosa* AmrZ (PA3385, alginate and motility regulator Z), was commonly enriched at an *e*-value of 5.2×10^−7^ (**Supplementary Table 2**). Collectively, these subset iModulons clearly indicate co-regulation by multiple TFs within the same reported regulon and provide a baseline for future regulation study for an in-depth understanding of associated TFs.

*ICA suggested an expanded regulatory role of Zur for metal uptake genes -* ICA identified the co-expression of 33 genes which were down-regulated when an excess amount of zinc was included in a medium or different carbon sources were utilized (**Fig. 2e** and **2f**). This iModulon, named Zur (MR of 0.52 and RR of 1.00, “regulon discovery”), included all seventeen genes (*znuBC*, *zur*, *znuA*, PP_0506-8, PP_1446, PP_3320-3, *folEB*, PP_5359-62), known to involve in zinc uptake and to be regulated by Zur (PP_0119)^39^. Interestingly, 16 new genes which do not have any reported regulation by Zur, were additionally detected by ICA to form an iModulon that is larger than the corresponding bottom-up defined regulon.

To investigate the possibility of common regulation by Zur, we enriched a common motif from the upstream sequence of all 19 TUs of the Zur iModulon genes. Notably, a common motif “WRRYGTTATRWTRTAACAWKWHWHW” was found in the upstream of 11 TUs including 22 genes (66.7%) at an *e*-value of 1.4×10^−27^ (**Fig. 2g**). This enriched motif is highly consistent with a previously known binding motif (TTGTTATAAGATAACAT) of Zur in *P. protegens* Pf-5 at an *E*-value of 1.39×10^−5^)^39,40^, supporting their regulation by Zur. The 22 genes include seven newly predicted target genes (PP_0509, PP_2416-9, PP_3325-6, **Fig. 2h**). In particular, the PP_2416-9 operon was annotated to encode an iron ABC transporter, which indicates that Zur has an expanded regulatory role in iron uptake, not limited to zinc. The activity changes depending on carbon sources (**Fig. 2f**), which may suggest different levels of metal requirements; further related studies are warranted. Nevertheless, these results show how iModulons in combination with binding motifs are useful to identify additional target genes of a given regulator.

*A ‘poorly-matched’ iModulon suggests conserved CysB regulation in P. putida -* We obtained a poorly-matched iModulon, PP_0204/CysB (originally named as PP_0204), which contains 29 genes in 17 TUs (**Fig. 2i,** MR of 0.10 and RR of 0.38). These iModulon genes were annotated to involve transportation of sulfur-containing compounds (e.g, *sbp-II*, *tauABC*, *ssuEADCB* encoding sulfate or sulfonate transporters) and showed high expression in the presence of hydrogen peroxide or glycolaldehyde, known to induce oxidative stress in microorganisms^25^ (**Fig. 2j**). Although PP_0204 (a GntR family TF) was originally enriched as a regulator, this TF was previously suggested to be a local regulator for itself and neighboring genes (e.g., PP_0205 and PP_0206 encoding an oxidoreductase and 4Fe-4S ferredoxin)^41^.

We hypothesized that there might be another major TF regulating these iModulon genes with a close interaction to PP_0204. This hypothesis was supported by a motif found from 11 of the total 17 TUs (22 genes of the 28 genes) at an *e*-value of 3.5×10^−21^ (**Fig. 2k**). To infer a potential main regulator of this iModulon, we compared its member gene weights with those of previously reported iModulons in *E. coli*^13^. Notably, the comparison showed that this iModulon is highly consistent with the Cbl+CysB iModulon in *E. coli*^42^ (**Fig. 2l**). Among the two *E. coli* TFs, we found that the *cysB* gene is also present in *P. putida* KT2440. In addition, CysB in the two strains shows high similarity (**Supplementary Fig. 6**), suggesting this TF is the main regulator. We further compared a previously reported binding motif of *E. coli* CysB with the enriched motif. Although the significance was relatively low (*p*-value of 3.86×10^−1^), the two motifs commonly share AT-rich sequences (**Fig. 2k**), also supporting common regulation for these genes by CysB in *P. putida*. Consequently, this iModulon was renamed as PP_0204/CysB based on these observations and this example shows that a poorly-matched iModulon can be evidence of regulation by an unstudied TF closely related to a linked TF.

### 2.3. iModulons provide a comprehensive description of transcriptome changes after the transition from the exponential growth phase to the stationary phase

Since iModulons represent biologically meaningful groups of co-regulated genes, complex transcriptomic changes can be simplified and comprehensively interpreted by monitoring iModulon activities rather than investigating the roles of huge numbers of differentially expressed genes (DEGs)^12,13^. To demonstrate this capability, we analyzed changed iModulon activities after the transition from the exponential growth phase to the stationary phase (project *phase*, **Supplementary Data**). When we applied the conventional DEG analysis, we obtained 1,454 genes (*p*_adj_<0.05) which correspond to 26.1% of the total number of genes (**Supplementary Fig. 7**).

In contrast, we obtained only 22 iModulons with differentially changed activities at an FDR < 0.05 and an activity threshold of 5; 67.8% of the total variance was explained by using iModulons (**Fig. 3a**). Among them, a total of sixteen iModulons showed increased activities while the activities of six iModulons were decreased. Changes of many iModulons were consistent with previous literature or reasonable in view of decreased metabolism and stress response due to nutrient depletion (**Supplementary Table 3**). For example, increased activities of PydR/RpoS for starvation responses, LiuR for ribose and lipid catabolism were identified whereas decreased activities of Translation for protein synthesis, and HexR for glucose metabolism were observed. Additionally, there were some iModulons (e.g., BkdR, Osmotic stress-1, GbdR, TurA-2) that showed unexpected activity changes; they can be targeted for detailed future studies.

**Figure 3.**
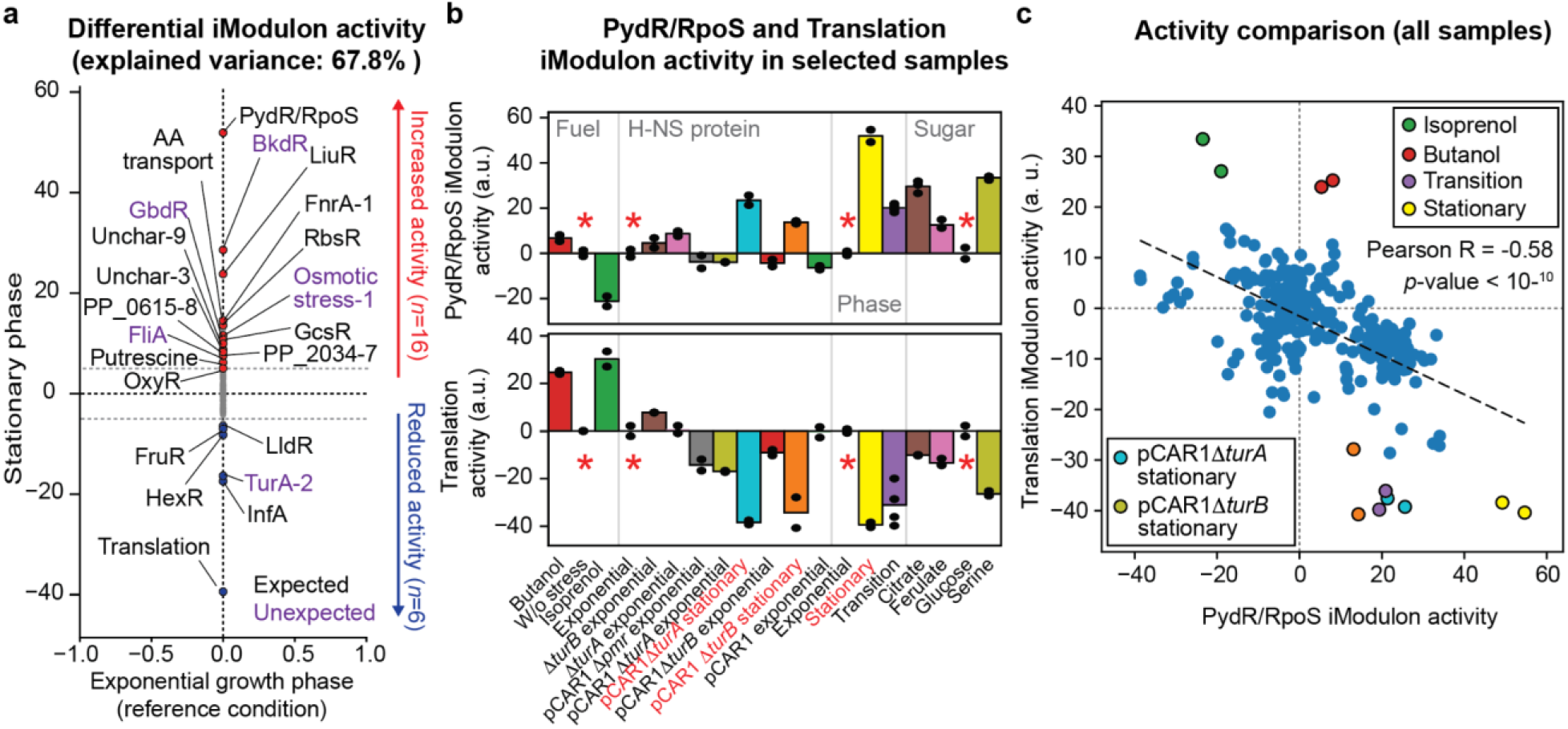
iModulons describe global transcriptional changes after the transition from the growth phase to the stationary phase. **(a)** Differentially changed iModulon activities during the transition from the exponential growth phase to the stationary phase at a threshold of 5 and an FDR of 0.05. Unexpected changes were colored in purple. (**b**) Activities of the PydR/RpoS and Translation iModulons in selected samples. Project names are colored in gray. Asterisks indicate the reference conditions for each project. (**c**) Activity correlation of the PydR/RpoS and Translation iModulons. Activity of these two iModulons showed a negative correlation across samples (Pearson correlation coefficient R of -0.58, *p*-value < 1×10^−10^).

The characteristics of PydR/RpoS and Translation iModulons were further investigated since they exhibited the greatest activity changes (**Supplementary Fig. 8a** and **8b**). PydR/RpoS, a regulatory iModulon (MR and RR of 0.17 and 1, respectively, regulon discovery), contains 30 genes presenting at various locations on the genome and explains the highest fraction of the total variance in our compendium (**Supplementary Fig. 3d**). Increased activities of the PydR/RpoS iModulon were consistently observed in all samples collected at the stationary phase in another project (*project*: H-NS protein, **Fig. 3b**). Its activity also changed depending on the carbon source (citrate, ferulate, or serine) or the presence of a stressor in the media. Originally, PydR was enriched as a major regulator; this TF is known to regulate the expression of four nucleobase degradation genes^43^ which display high gene weights in this iModulon: *pydXA*, *hyuC*, and *pydB* (**Supplementary Fig. 8a**). However, additional member genes for carbohydrate transporters (e.g., PP_1121), stress-mitigation (e.g., *katG* encoding catalase peroxidase and PP_1121 encoding OmpA family protein), and energy generation (e.g., *icd* encoding NADP^+^-specific isocitrate dehydrogenase) were identified. Notably, a motif was enriched from all 27 TUs, including the PydR regulon genes, at an *e*-value of 2.8 × 10^−7^ (**Supplementary Table 2**), suggesting that not only are the four genes regulated by a common regulator, but all the other genes are as well. In many microorganisms including *E. coli*, a similar stationary response was shown to be governed by RpoS and a RpoS iModulon explained the largest variance in a corresponding gene expression compendium^11–13^. Furthermore, its activity also showed a high correlation with the expression level of *rpoS* rather than *pydR* (**Supplementary Fig. 9a** and **9b**). Collectively, this iModulon may represent the regulation by RpoS (PP_1623) in *P. putida* and thus renamed to PydR/RpoS.

On the other hand, the Translation iModulon, a functional iModulon with an unknown regulator, includes 45 genes that are mostly ribosomal proteins and clustered together on the genome (**Supplementary Fig. 8b**). Additionally, it includes genes encoding RNA polymerases (*rpoB* and *rpoC*), elongation factors (e.g., *tufA*, *fusA*), cytochrome oxidases (*cyoC* and *cyoD*), and ATP synthase subunits (*atpG* and *atpA*), important for protein synthesis and energy production. Therefore, its reduced activity at the stationary phase indicates decreased biomass production. Interestingly, a global inverse correlation of the PydR and Translation iModulon activities were observed across samples regardless of growth phases (Pearson coefficient of -0.58 at a *p*-value < 10^−10^, **Fig. 3c**), except a few conditions (e.g., butanol addition).

Previously, a similar negative correlation between stationary-phase induced genes and growth-related genes was described as a major transcriptomic trade-off in *E. coli*^13^. The observation in *P. putida* indicates a similar trade-off is conserved across species for diverting resources from growth to maintenance. Collectively, the changed iModulon activities clearly illustrate the transcriptional changes that occurred during the transition from the growth phase to the stationary phase.

### 2.4 iModulons identify substrate-specific gene groups and show increased stationary phase response during utilization of poor carbon sources

We examined how iModulon activities reflect the utilization of different substrates. Although catabolic efficiencies vary depending on substrates, resulting in global changes in growth rates, understanding of transcriptome changes in *P. putida* depending on substrates has been limited. *putida*PRECISE321 includes seventeen gene expression profiles with 11 different carbon source conditions: glucose (as a reference), gluconate, fructose, galactose, xylose, citrate, coumarate, ferulate, serine, acetate, and myristic acid. In addition, it should be noted that gene expression profiles with xylose and galactose utilization were obtained from engineered *P. putida* strains with heterologous gene expression^26^.

ICA identified multiple iModulons (e.g., HexR, GltR-II, FruR, PsrA, PP_5350) containing genes responsible for transportation and catabolism for each substrate (**Fig. 4a**). Except for the GlnG iModulon, the activity of most iModulons specifically showed increased or decreased activity depending on catabolic routes for each carbon source (**Fig. 4b**). For example, the PsrA and PP_5350 iModulons responsible for β-oxidation and glyoxylate shunt pathway, respectively, showed high activities only when myristic acid, a C-14 fatty acid, was utilized as a sole carbon source. These observations indicate that the expression of catabolic genes is tightly regulated by multiple related TFs. Additionally, high activities of the GlnG iModulon were observed in most non-glucose conditions (e.g., galactose, xylose, fructose) in addition to the serine utilization condition. This iModulon is the regulon subset of GlnG (PP_5048, also called NtrC) is known to control the expression of 63 nitrogen metabolism genes^44^. Although it is not clear, this observation implied that intracellular nitrogen availability could be affected depending on carbon sources.

**Figure 4.**
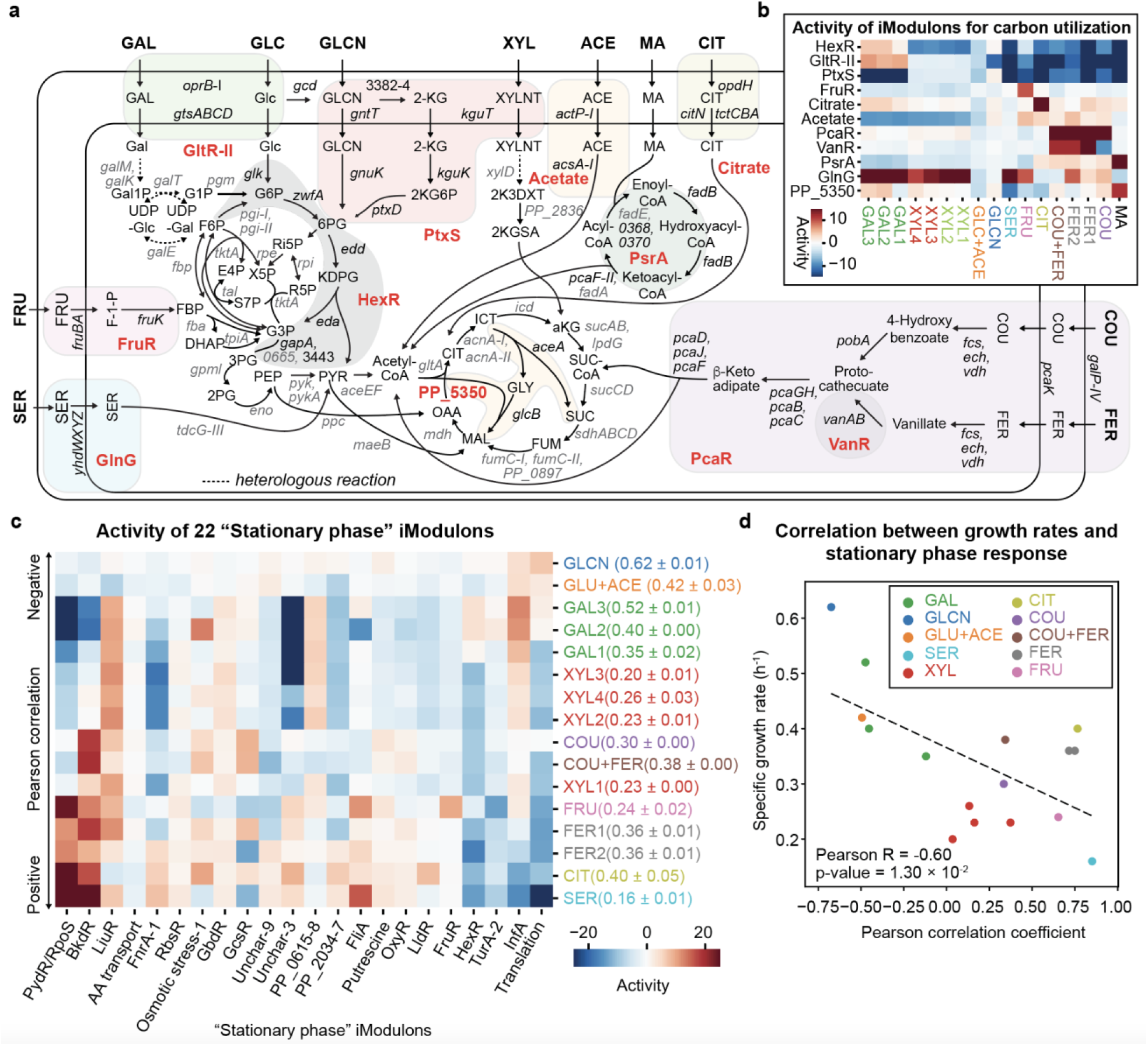
iModulons show transcriptional responses to the utilization of different substrates. (**a**) Metabolic pathway associated with iModulons for substrate utilization in *P. putida*. Boundaries of iModulons (names in red) were indicated by colored rounded shapes. Names of genes belonging to iModulons were colored in black. Subcellular localization information was obtained from Pseudomonas.com^64^. (**b**) Activities of the carbon utilization iModulons depending on substrates. For a clear comparison, the minimum and maximum activity values were set to -15 and 15, respectively. (**c**) The activities of the 22 iModulons, that showed differential activities during the transition to the stationary phase from the growth phase, during utilization of each substrate. For clear visualization, the minimum and maximum activities were set to -25 and 25, respectively. (**b** and **c**) Multiple RNA-Seq samples with the same substrates were differentiated by adding numbers. iModulon activities in the glucose condition are zero. (**d**) A scatter plot of the growth rates and Pearson correlation coefficients of activities of the “stationary response” in iModulons in the group. Abbreviations: GAL, galactose; GLC, glucose; GLCN, gluconate; XYL, xylose; ACE, acetate; MA, myristic acid; CIT, citrate; FRU, fructose; SER, serine; 2-KG, 2-ketogluconate; XYLNT, xylonate; 2K3DXT, 2-keto-3-deoxyxylonate; 2KGSA, 2-keto-glutaric semialdehyde; G6P, glucose-6-phosphate (P); 6PG, 6-phosphogluconate; 2KG6P, 2-ketogluconate-6-P; Gal1P, galactose-1-P; F-1-P, fructose-1-P; UDP-Glc, uridine diphosphate-glucose; UDP-Gal, uridine diphosphate galactose; G1P, glucose-1-P; KDPG, 2-dehydro-3-deoxy-phosphogluconate; PYR, pyruvate; G3P, glyceraldehyde-3-P; 3PG, glycerate-3-P; 2PG, 2-phosphoglycerate; PEP, phosphoenolpyruvate; acetyl-CoA, acetyl coenzyme A (CoA), CIT, citrate; ICT, isocitrate; αKG, α-ketoglutarate; SUC-CoA, succinyl-CoA; SUC, succinate; FUM, fumarate; MAL, malate; OAA, oxaloacetate; GLY, glyoxylate

We further hypothesized that poor carbon sources, which result in low growth rates, induce the stationary phase-like response in the transcriptome even if the culture is still in the exponential growth phase. To test this hypothesis, we investigated the activity of the “stationary phase” iModulons (i.e., the 22 iModulons that showed differential activity during the transition from the growth phase to the stationary phase) with specific growth rates (0.16 h^−1^ to 0.62 h^−1^, **Fig. 4c** and **4d**). Notably, we observed that the activity of the 22 iModulons in cells growing on gluconate, a mixture of glucose and acetate, and galactose clearly showed a negative correlation (up to a Pearson correlation of -0.83 at a *p*-value of 8.54 × 10^−4^) whereas cells growing on xylose, aromatic compounds (i.e., coumarate and ferulate), citrate, and serine showed positively correlated iModulon activity (up to a Pearson correlation of 0.89 at a *p*-value of 0.96 × 10^−4^). We found that the correlation of a sample’s activity levels with stationary phase activity levels is itself correlated with growth rate, indicating that stationary phase iModulon activities are a fairly conserved transcriptomic signature for slowed growth. This correlation could be utilized to estimate growth rates from a given transcriptome. Taken together, these results show that global correlations in transcriptomes can be efficiently found by analyzing iModulon activities in multiple datasets, revealing tradeoffs in transcriptome composition.

### 2.5. iModulons unveil evolutionary reallocation of the transcriptome to achieve fast growth rates

Once an iModulon structure for an organism is established, it can be used to interpret new data sets by inferring the activities of each iModulon based on the gene weights from the [**M**] matrix of the original compendium. Thus, we examined the utility of the *P. putida* iModulon structure to unveil an evolutionary strategy for achieving high fitness in a glucose-minimal medium. We analyzed iModulon activity in three evolved *P. putida* KT2440 strains (A1_F11_I1, A3_F85_I1, and A4_F88_I1, see **Supplementary Note 3** for naming)^24^, displaying higher growth rates (0.64 h^−1^, 0.77 h^−1^, and 0.76 h^−1^, respectively) over the wildtype (0.58 h^−1^, **Fig. 5a** and **5b**). These strains were independently evolved but all acquired mutations in *relA*, which encodes one of two guanosine tetraphosphate (ppGpp) synthetases in *P. putida*, suggesting its critical role in the improved fitness. The A1_F11_I1 strain which was isolated from an earlier flask (11^th^) in a laboratory evolution study, only has a single mutation - a frame-shift in *relA*, whereas the two latter strains have different additional mutations in other genes in addition to a single amino acid change or early termination mutation in *relA*.

**Figure 5.**
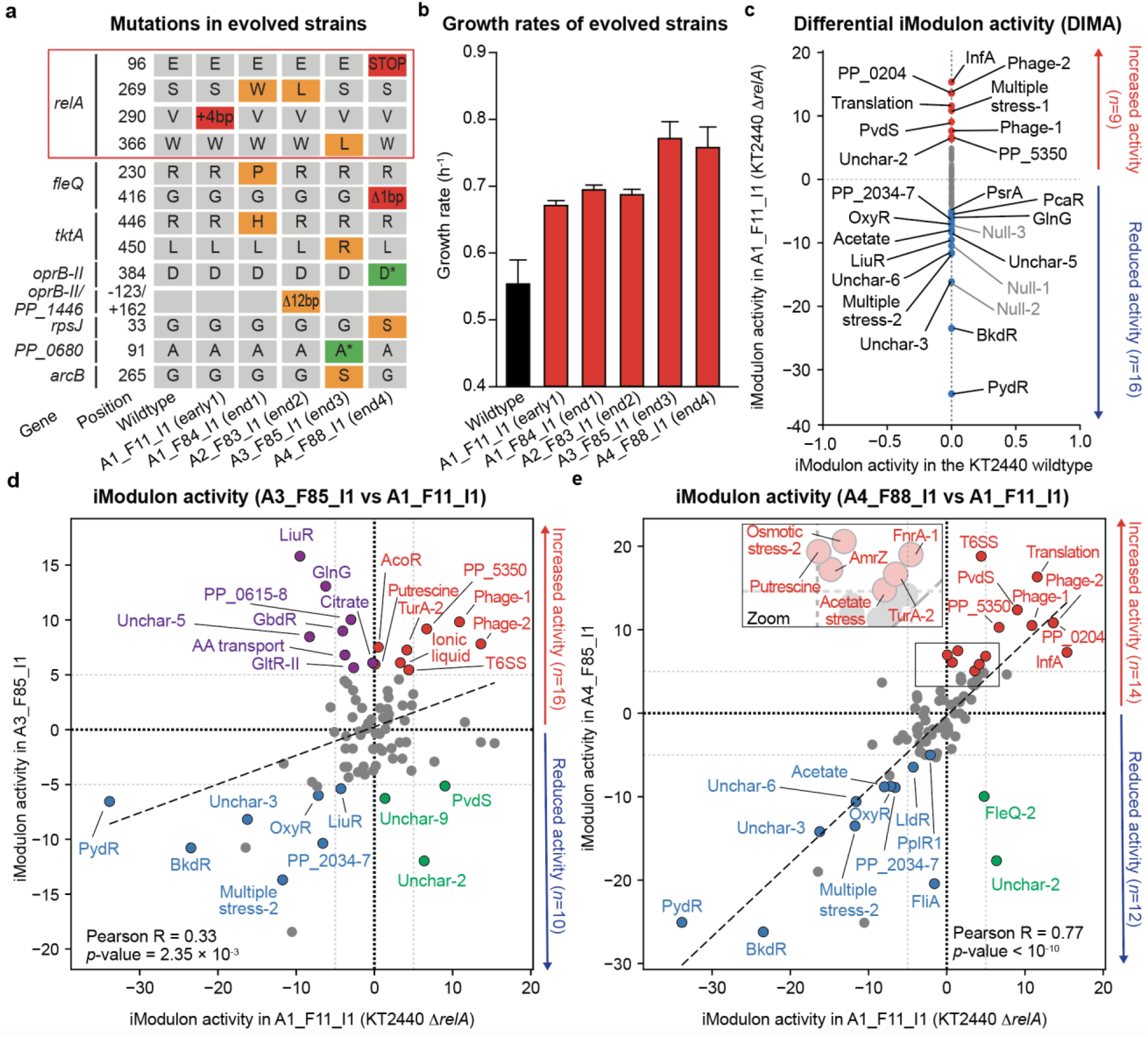
Analyzing iModulon activities unveil evolutionary strategies to achieve high fitness in a glucose minimal medium condition. (**a**) Mutations identified in the wildtype, A1_F11_I1 (early1), A1_F84_I1 (end1), A2_F83_I1 (end2), A3_F85_I1 (end3), and A4_F88_I1 (end4) strains^24^. Strain names indicate passage numbers during an evolution study (see **Supplementary Note 3**). Colors: orange, single amino acid changes or an in-frame deletion; green, nonsynonymous mutation; red, early termination or frame-shift mutations (**b**) Maximum specific rates (h^−1^) of the six strains in a glucose minimal medium. (**c**) Differential iModulon activity plot for the A1_F11_I1 and wildtype strains. (**d** and **e**) Differential iModulon activity plot for the A1_F11_I1 versus either (**d**) A3_F85_I1 or (**e**) A4_F88_I1 strain. iModulon activities in the non-wild type strains were inferred from the [**M**] matrix of the original iModulon activity. Dashed gray lines indicate a threshold value of 5 and solid gray lines indicate a 45-degree line of each plot. Null iModulons were colored in gray even with changes greater than 5.

Initially, we investigated changes in iModulon activities in the A1_F11_I1 strain (KT2440 with the loss-of-function mutation in *relA*) to understand the effect of reduced ppGpp synthesis (**Fig. 5c**). We found 22 iModulons with large activity changes (using a threshold of 5). Nine iModulons including Translation and InfA responsible for cell growth showed increased activities whereas thirteen iModulons including starvation iModulons (PydR and BkdR) and stress-hedging iModulons (OxyR, Multiple stress-2) showed decreased activities. These observations were consistent with previous studies that ppGpp has a pivotal role to determine whether cells actively grow or show quiescence phenotypes^45^. In addition, it also shows efficient reallocation of cellular resources to obtain robust cell growths in the evolution condition.

We further compared iModulon activities in the A3_F85_I1 and A4_F88_I1 strains against the A1_F11_I1 strain (**Fig. 5d** and **5e**). These two strains showed even higher growth rates (0.77 h^−1^ and 0.76 h^−1^) over the A1_F11_I1 strain. However, no genes other than *relA* were commonly mutated in these strains, suggesting different evolutionary strategies for improving growth rates. When the two strains were compared with the A1_F11_I1 strain, their transcriptomes have an overall positive correlation (Pearson R values of 0.33 and 0.77 at *p*-values of 2.35 × 10^−3^ and less than 10^−10^, respectively), likely owing to mutations commonly occurred in *relA*. Interestingly, we identified several iModulons with inversely correlated activity changes in the two strains. In the A3_F85_I1 strain, eight iModulons (colored in purple in **Fig. 5d**, e.g., LiuR, GlnG, PP_0615) showed high activities, whereas three iModulon (Uncharacterized-2, Uncharacterized-3, PvdS, colored in green) showed decreased activities. On the other hand, in the A4_F88_I1 strain, only two iModulons (FleQ-2 and Uncharacterized-2, colored in green in **Fig. 5e**) showed decreased activity.

From these iModulon activity changes, two distinctive evolution strategies could be inferred. In the A3_F85_I1 strain, three mutations in *tktA* (a gene encoding transketolase A), PP_0680 (a gene encoding ATP-dependent protease-like protein), and *arcB* (a gene encoding ornithine carbamoyltransferase) likely activated nutrient re-assimilation or utilization; their effects were shown as increased activities of iModulons for Ribose and lipid catabolism (LiuR), amino acid metabolism (GlnG, AA transport), carbon metabolism (GltR-II). Differently, the decreased activity of FleQ-2 containing flagella synthesis genes implies that the A4_F88_I1 strain achieved efficient biomass formation by minimizing wasteful synthesis of extracellular proteins from the frame-shift mutation in *fleQ*^46^. Taken together, multiple evolutionary strategies were identified by analyzing iModulon activities. This method has significant potential to be broadly applicable to identify meaningful information from new gene expression profiles.

### 2.6. Coordinated iModulon activity changes define a stimulon and underlie the activation of the Type VI secretion system (T6SS) by osmotic stress

Activities of iModulons may be correlated, especially if they share master regulators or common activating conditions. Therefore, clustering the activity levels can elucidate underlying relationships or hierarchical transcriptional regulation and define a stimulon (i.e., genes with changed expression by an external stimulus^47^). As an example, we found that activities of four iModulons (Osmotic stress-1, Osmotic stress-2, AmrZ, T6SS) showed high correlations across samples (**Fig. 6a**). The Osmotic stress-1 and Osmotic stress-2 iModulons included 46 and 31 genes, respectively, commonly showing high activities under osmotic-stressed conditions but displaying different strengths depending on the types of osmotic stress (i.e., NaCl or protic ionic liquid addition). The AmrZ iModulon included 12 genes for alginate synthesis. The high correlation of the first three iModulons recapitulates previous findings that *P. putida* increases the synthesis of alginate, which is one of the biofilm components, to create a microhydrated environment under high-osmotic stress conditions^48,49^. Meanwhile, the T6SS iModulon includes 15 machinery genes for the Type VI secretion system (T6SS, **Fig. 6b**) required for its antibacterial activity against phytopathogens (e.g., *Xanthomonas campestris*), which functions by injecting toxins^50^. In particular, this iModulon showed high activities in samples collected under regrowth phase (i.e., promoting cell growth by additional supplementation of a carbon source after its depletion) and NaCl-treated samples (**Fig. 6c**). Given that the T6SS iModulon was clustered together, it can be suggested that the secretion system is induced by osmotic changes. Indeed, a close relationship between the activity of T6SS and biofilm formation was reported in *P. aeruginosa*^51^ and the activation of T6SS by OmpR, a global osmostress regulator, was previously suggested for *Klebsiella pneumoniae*^52^. Overall, our iModulon clustering provides strong evidence that the T6SS system is activated by osmotic changes in a surrounding environment, and reflects the classical definition of a stimulon.

**Figure 6.**
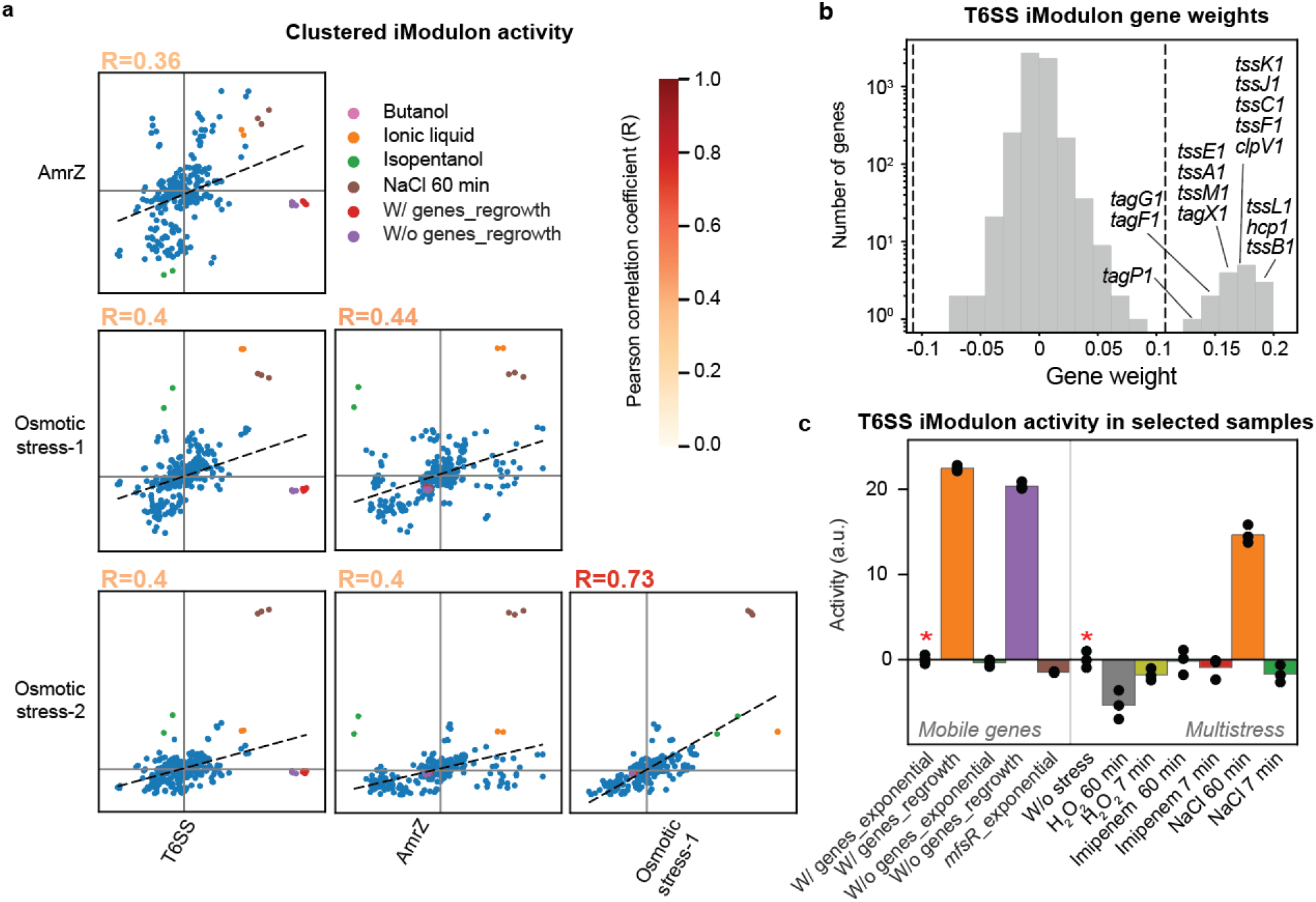
Coordinated iModulon activity changes underlie the activation of the Type VI secretion system by osmotic stress and define a stimulon. (**a**) Activity correlation of the T6SS, Osmotic stress-1, Osmotic stress-2, and AmrZ iModulons. Activities of all these four iModulons showed a positive correlation. Font colors indicate Pearson correlation coefficients. Samples displaying high activities were colored: pink (fuel:butanol); green (fuel:isopentanol), orange (ALE:IL_TEA_HS), brown (multistress:NaCl_60min), red (mobile_genes:2737_reg-w/ mobile genes at the regrowth phase), and purple (mobile_genes:3227_reg-w/ mobile genes at the regrowth phase). Dashed lines are the best fits on each plot. (**b**) Gene weights of the T6SS iModulon. The dashed line indicates a threshold value of 0.11 (See Methods). (**c**) T6SS iModulon activities in selected samples from projects: *mobile genes*^23^ and *multistress*^25^. See Supplementary Data for details of the experimental conditions. Asterisks indicate the reference conditions.

## 3. Discussion

In this study, we uncovered the TRN of *P. putida* by obtaining 84 iModulons using ICA of 321 high-quality RNA-seq samples. 39 Regulatory iModulons with linked TFs, recapitulated previously reported TF-gene interactions and identified knowledge gaps (e.g., the existence of a potential co-regulator or a new regulation target) via a systematic comparison of iModulon gene membership and existing TRNs. In addition, we identified 24 groups of genes as functional iModulons and suggested them as excellent targets for future studies to elucidate unreported regulators, given their co-expression upon environmental changes (e.g., osmotic stress). Compared to traditional regulon discovery approaches which investigate activities in a few experimental conditions, ICA simultaneously analyzes various gene expression profiles and identifies multiple gene groups likely under the same regulation in a high-throughput manner.

Since ICA provides groups of genes, regulators can be efficiently discovered when combined with motif search and comparison using related tools such as MEME^35^ and TOMTOM^38^. We found 32 common motifs in the upstream of TUs in each iModulon (**Supplementary Table 2**), which strongly supports co-regulation by a common regulator(s). We showed that the common regulators can be inferred by comparing them with previously known motifs across species. Out of the 32 motifs, 8 motifs from the FleQ/Fur, FnrA-2, HexR, Multiple stress-1, PvdS, PydR/RpoS, TCA cycle, Zur iModulons showed similarity to known TF binding motifs. Therefore, iModulon and motif analysis can be combined to identify a regulator(s) and cross-species comparison of TRNs.

iModulons are powerful to visualize key changes in gene expression profiles. In contrast, DEG analysis is not suitable for the identification of global changes in multiple gene expression profiles containing thousands of genes. Although the 26 categories of Clusters of Orthologous Groups of proteins (COGs)^53^ are often utilized, the category size is relatively small, and such categories do not consider individual characteristics of a specific microorganism of interest. Conversely, as shown in this work, iModulons were defined from gene expression profiles of a given microorganism and thus they are specific to the corresponding microorganism. Moreover, the number of iModulons is determined after a screening of the optimal dimensionality to avoid both under-decomposition and over-decomposition of a gene expression compendium^54^. With the optimum reduction in complexity, various transcriptomes, even newly generated, can be comprehensively interpreted.

By using iModulons, we comprehensively explained transcriptome changes in various conditions that are closely related to industry-relevant bioprocesses (i.e., carbon-depletion induced stationary phase, utilization of different carbon sources with different catabolic efficiencies, evolutionary adaptation to a glucose minimal medium condition). Although understanding the stationary phase response is essential, its understanding at a glance was limited owing to an absence of tools. In this regard, we identified 22 iModulons with differentially changed activities after transitioning to the stationary phase from the exponential growth phase. These iModulons indicate how *P. putida* reallocates its transcriptome under a nutrient-depleted environment for its survival. Moreover, it was shown that slow growth rates on poor substrates induce the stationary phase-like response even at the exponential growth phase. Finally, multiple evolutionary strategies to achieve high growth rates were unveiled by using iModulons. iModulon-based interpretation of transcriptome changes that occurred during diverse bioprocesses will aim a better understanding of cellular responses and subsequent strain designing.

The iModulon results can be utilized in a variety of future studies. They can be subjected to detailed experimental validation via RNA-seq of TF-deleted strains or ChIP-sequencing^10^ to confirm suggested TF-gene relationships. Regulators for functional or uncharacterized iModulons could be identified by using the motif information via DNA-pull down assay. In addition, since iModulons allow efficient comparison across multiple conditions, they can generate many hypotheses from the activity comparison. In particular, based on the activity clustering, we suggested that the activity of the T6SS is closely related to osmotically stressed conditions by monitoring gene expressions in samples. In a similar vein, new hypotheses can be generated and subsequently tested to expand our understanding of *P. putida*. The functions of uncharacterized genes (i.e., unknown function genes) can be also inferred from active conditions. Additionally, since iModulons provide boundaries of genes required for a given functionality (e.g., aromatics catabolism) with statistical significance, identified gene sets can be studied for reinforcement of functions or even heterologously transferred to another microorganism for engineering.

Lastly, the explanatory power of iModulons can be further increased by performing ICA for a larger scoped compendium to achieve a more detailed structure of the TRN. For example, the second round of ICA was performed for 815 *E. coli* RNA-seq samples^42^ to identify 218 iModulons, which were expanded from 92 iModulons reported in the first version^13^; the expansion covering diverse genetic perturbations and environmental conditions generated more detailed iModulon structures and new regulon discoveries^42^. Efforts can be extended for *P. putida* to identify a more detailed iModulon structure underlying its TRN.

Taken together, this study provides the first global TRN analysis for *P. putida*, and we believe that the *P. putida* iModulons will be useful for advancing our fundamental understanding and engineering of this microorganism for many different purposes.

## 4. Methods

### 4.1 Generation of a high-quality *P. putida* RNA-Seq compendium

To perform ICA, a total of 541 *P. putida* RNA-Seq samples were collected; 435 samples were obtained from the NCBI Sequence Read Archive (SRA, https://www.ncbi.nlm.nih.gov/sra, published before Aug 31, 2020); 90 samples were obtained from a previous study^22^; 16 samples were newly generated in this study (**Supplementary Data**). RNA-seq samples, generated in this study, were prepared as described in **Supplementary Method 1**. Raw sequencing files were processed by the prokaryotic RNA-seq processing pipeline^18^ (https://github.com/avsastry/modulome-workflow) implemented to use Nextflow v20.01.0^55^ on Amazon Web Services (AWS) as previously described^18^. Briefly, the pipeline utilizes fasterq-dump (https://github.com/ncbi/sra-tools/wiki/HowTo:-fasterq-dump), Trim Galore (https://www.bioinformatics.babraham.ac.uk/projects/trim_galore), FastQC (http://www.bioinformatics.babraham.ac.uk/projects/fastqc/) for raw data processing. Sequencing reads were aligned to the reference genome (AE015451.2) using Bowtie^56^. Read counts were generated by using RSEQC^57^ and featureCounts^58^ and converted into log_2_ transcripts per million (TPM).

To ensure the quality of the compendium (**Supplementary Fig. 1**), MultiQC^59^ was performed and failed samples were discarded based on per base sequence quality, per sequence quality scores, per base n content, and adapter content. Hierarchical clustering was also utilized to identify and discard datasets not displaying a typical expression profile. Furthermore, metadata for each sample was manually curated by pulling information from literature and the registry and samples with low correlation within biological replicates (R^2^ < 0.95) were discarded. Additionally, non-*P. putida* KT2440 samples were excluded for consistency of gene expression profiles. For 321 samples that passed the aforementioned QC metrics, unique project and condition identifiers were given (*project:condition*, **Supplementary Data**).

### 4.2 Independent component analysis (ICA)

ICA^16,18^ was performed as introduced in previous studies^13,18^ to decompose the gene expression profile [**X**] into independent components (i.e., iModulons, [**M**]) and their activities [**A**] in each sample.

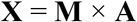

Briefly, each gene expression profile was normalized by dividing the log TPM values by the values of a reference condition within each project (**Supplementary Data**) to remove potential batch effects. Then, the scikit-learn (v0.23.2)^60^ implementation of FastICA (Hyvärinen, 1999) was performed 100 times with a convergence tolerance of 10^−7^ to obtain robust independent components (ICs). The resulting ICs were clustered by using DBSCAN (Ester et al., 1996) to identify robust ICs from each random restart. For the clustering, an epsilon of 0.1 and a minimum cluster seed size of 50 were used. Because identical ICs can have opposite signs, distance (*d*) between components was calculated from the absolute value of the Pearson correlation (*ρ)* as follows:

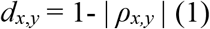

To choose the optimal dimension^54^, we performed the clustering to the gene expression profile multiple times for dimensions between 10 to 290 with the step size of 10. The optimal dimension of 220 was chosen where the number of non-single gene ICs was equal to the number of final components. Finally, the **M** and **A** matrices for 84 robust ICs (i.e., iModulons) were obtained. From the obtained [**M**] matrix, iModulon member genes for each iModulon were determined by choosing an optimal threshold, as explained in **Supplementary Method 2**.

### 4.3 Characterization of iModulons

The 84 iModulons were characterized by the Pymodulon package (https://github.com/SBRG/pymodulon)^18^. Functions of the iModulons were inferred by enriching known regulators, Kyoto Encyclopedia of Genes and Genomes (KEGG, https://www.genome.jp/kegg/) annotations of member genes. In particular, we collected 1993 TF-gene interactions (i.e., regulon) in *P. putida* (and some in *P. aeruginosa*) from RegPrecise 3.0^61^, BioCyc Database Collection^62^, and literature. All interactions and their sources are listed in **Supplementary Data**. Subsequently, Fisher’s Exact Test was performed with an FDR of < 10^−4^ using the Benjamini-Hochberg correction. KEGG annotations and Cluster of Orthologous Groups (COG) information were obtained by running eggNOG-mapper (version 5)^63^. Fisher’s Exact test was also utilized to enrich GO and KEGG annotations with an FDR of < 10^−2^. Common motifs in upstreams of iModulons were enriched by using the find_motifs function that uses MEME^35^ with a maximum window of 30 bp. When motifs were compared, the web-based TOMTOM^38^ was utilized. As a statistical significance, *E*-values were provided as adjusted *p*-values when more than one common sequence was identified or compared. iModulons were named with their enriched TFs from the regulons or functional characteristics. When additional TF was suggested from a motif search or a comparison of iModulons in other microorganisms, iModulon names were changed accordingly. Additionally, agglomerative activity clustering of iModulons was performed by using the cluster_activities function. iModulon activities in new expression profiles were inferred by using the infer_activities function.

### 4.4. Generating iModulonDB web page

The *P. putida* iModulonDB^21^ page was generated by using the imodulondb_export function in the Pymodulon package^18^ and it is accessible at https://imodulondb.org. The generated page includes gene information from the Pseudomonas Genome DB (https://pseudomonas.com)^64^.

## Data availability

All RNA-seq raw files are available at GEO with accession numbers listed in Supplementary Data. Source codes for iModulon analysis and figures are available at https://github.com/SBRG/modulome_ppu.

## Acknowledgments

This work conducted by the Joint BioEnergy Institute was supported by the Office of Science, Office of Biological and Environmental Research, of the U.S. Department of Energy under Contract No. DE-AC02-05CH11231. This work was also supported by the Nebraska Center for Energy Science Research at the University of Nebraska - Lincoln (Cycle-11 grant to W.N.). We also thank Richard Szubin, Ying Hutchison, and Marc K. Abrams for experimental support and manuscript editing.

## Author contributions

H.G.L. and B.O.P. conceived the project. H.G.L. collected and analyzed the data, together with K.R. and A.V.S. J.M. generated RNA-seq libraries for the aromatic compound utilization condition with the guidance of W.N. All authors participated in writing the manuscript. B.O.P. supervised the project. All authors read and approved the final version of the manuscript.

## Supplementary information

### Supplementary Notes

Supplementary Note 1. Summary of 84 iModulons

Supplementary Note 2. Nucleotide symbols

Supplementary Note 3. The naming of evolved strains

### Supplementary Methods

Supplementary Method 1. Transcriptome sequencing (RNA-seq)

Supplementary Method 2. Determination of iModulon threshold values

### Supplementary Tables

Supplementary Table 1. Selected 30 Regulatory iModulons

Supplementary Table 2. The best enriched motif from transcriptional units in iModulons

Supplementary Table 3. iModulons with differential activity changes between the exponential growth phase and the stationary phase

### Supplementary Figures

Supplementary Figure 1. General information of collected RNA-seq samples

Supplementary Figure 2. Principal component analysis of putidaPRECISE321

Supplementary Figure 3. Characteristics of the 84 P. putida iModulons

Supplementary Figure 4. Types of 39 regulatory iModulons depending on iModulon and Regulon recalls

Supplementary Figure 5. Characteristics of the two FleQ-related iModulons (FleQ/Fur and FleQ/AmrZ)

Supplementary Figure 6. Comparison of CysB from E. coli K-12 MG1655 and P. putida KT2440

Supplementary Figure 7. Differentially expressed gene analysis of the transcriptomes at the exponential and stationary phase

Supplementary Figure 8. Gene weights of the PydR/RpoS and Translation iModulons

Supplementary Figure 9. Correlation between the activity of PydR/RpoS iModulon and the expression level of either pydR and rpoS

### Supplementary Data

Supplementary Data (xlsx) - all information is available at: https://github.com/SBRG/modulome_ppu.

Sheet 1: RNAseq sample list

Sheet 2: Gene table

Sheet 3: TRN

Sheet 4: iModulon table

Sheet 5: The X matrix

Sheet 6: The M matrix

Sheet 7: The A matrix

