## Supplementary information for "Machine-learning from *Pseudomonas putida* Transcriptomes Reveals Its Transcriptional Regulatory Network"

(Bernhard O. Palsson)

Mailing address: Department of Bioengineering, University of California San Diego, 9500 Gilman Drive La Jolla, CA, USA, 92093-0412

### Supplementary Notes

### Supplementary Note 1. Summary of 84 iModulons

All detailed iModulon information including member gene weights, active conditions is available at https://iModulonDB.org[1]. Pymodulon source code for *P. putida* is available at: https://github.com/SBRG/modulome_ppu.

#### 1. PydR/RpoS iModulon (*n*=30, explained variance: 5.4%)

Category: Regulatory iModulon (MR and RR of 0.17 and 1.00, poorly-matched)

Inferred role: stationary or starvation response

Related regulators: PydR (PP_4039), RpoS (PP_1623)

Top five genes with highest weights:

| Locus tag | Gene weight | Gene name | Gene product |
|---|---|---|---|
| PP_4037 | 0.098686 | *pydX* | NADP-dependent dihydropyrimidine dehydrogenase subunit |
| PP_4038 | 0.096952 | *pydA* | NADP-dependent dihydropyrimidine dehydrogenase subunit PreA |
| PP_0679 | 0.094899 | *PP_0679* | conserved exported protein of unknown function |
| PP_1121 | 0.093217 | *PP_1121* | OmpA family protein |
| PP_4034 | 0.091995 | *hyuC* | N-carbamoyl-beta-alanine amidohydrolase/allantoine amidohydrolase 2 |

Differentially active condition: stationary phase

#### 2. Genomic-1 (*n*=19, explained variance: 4.9%)

Category: Genomic iModulon

Active condition: Muconate study (deletion)

#### 3. TurA-1 iModulon (*n*=17, explained variance: 4.0%)

Category: Regulatory iModulon (MR and RR of 0.94 and 0.07, regulon subset)

Inferred role: Biosynthesis of lipodepsinonapeptide phytotoxins

Regulator: TurA (PP_1366)

Top five genes with highest weights:

| Locus tag | Gene weight | Gene name | Gene product |
|---|---|---|---|
| PP_3783 | 0.279933 | *syrB* | syringomycin biosynthesis enzyme 2 |
| PP_3781 | 0.27732 | *PP_3781* | Oxygen-independent Coproporphyrinogen III oxidase family protein |
| PP_3782 | 0.275988 | *PP_3782* | conserved protein of unknown function |
| PP_3784 | 0.24789 | *PP_3784* | conserved protein of unknown function |
| PP_3785 | 0.236145 | *PP_3785* | conserved protein of unknown function |

Differentially active condition: Deletion of *crc*, *crcZY*, or *turA*

#### 4. Multiple stress-1 iModulon (*n*=58, explained variance: 3.4%)

Category: Functional iModulon

Inferred role: arginine metabolism, oxygen or nitrogen limitation

Potential regulator: FnrA (PP_4265, also known as Anr), FnrB (PP_3233), FnrC (PP_3287)

Top five genes with highest weights:

| Locus tag | Gene weight | Gene name | Gene product |
|---|---|---|---|
| PP_1002 | 0.133642 | *arcD-I* | arginine/ornithine antiporter |
| PP_5591 | 0.113572 | *PP_5591* | conserved protein of unknown function |
| PP_0807 | 0.110851 | *norR* | DNA-binding transcriptional dual regulator (NO) |
| PP_3931 | 0.104498 | *yfbS* | putative transporter |
| PP_1001 | 0.101448 | *arcA* | arginine deiminase |

Differentially active condition: substrate type

#### 5. BkdR iModulon (*n*=19, explained variance: 2.8%)

Category: Regulatory iModulon (MR and RR of 0.16 and 0.75, regulon discovery)

Inferred function: starvation response

Regulator: BkdR (PP_4400)

Top five genes with highest weights:

| Locus tag | Gene weight | Gene name | Gene product |
|---|---|---|---|
| PP_0105 | 0.10509 | *PP_0105* | Cytochrome C oxidase assembly protein |
| PP_1640 | 0.102939 | *yohC* | putative inner membrane protein of unknown function |
| PP_2361 | 0.090849 | *PP_2361* | putative chaperone protein |
| PP_2359 | 0.090422 | *PP_2359* | putative Type 1 pili subunit CsuA/B protein |
| PP_2360 | 0.09013 | *PP_2360* | putative type 1 pili subunit CsuA/B protein |

Differentially active condition: carbon sources, growth state, oxidative stress

#### 6. PedR1-2 iModulon (*n*=31, explained variance: 2.6%)

Category: Regulatory iModulon (MR and RR of 0.84 and 0.76, well-matched)

Inferred function: alcohol/aromatics catabolism

Regulator: PedR1 (PP_2665)

Top five genes with highest weights:

| Locus tag | Gene weight | Gene name | Gene product |
|---|---|---|---|
| PP_2675 | 0.26206 | *PP_2675* | Cytochrome c-type protein |
| PP_2676 | 0.256452 | *PP_2676* | putative Periplasmic binding protein |
| PP_2677 | 0.24344 | *PP_2677* | conserved exported protein of unknown function |
| PP_2678 | 0.23939 | *PP_2678* | putative Hydrolase |
| PP_2669 | 0.228156 | *PP_2669* | putative Outer membrane protein |

Differentially active condition: carbon sources (ferulic acid), oxidative stress, antibiotic stress,

### 7. Translation iModulon (*n*=46, explained variance: 2.4%)

Category: Functional iModulon

Inferred function: Protein synthesis

Regulator: unknown

Top five genes with highest weights:

| Locus tag | Gene weight | Gene name | Gene product |
|---|---|---|---|
| PP_0456 | 0.108925 | *rplW* | 50S ribosomal protein L23 |
| PP_0451 | 0.108097 | *fusA* | Elongation factor G 1 |

| | | | |
|---|---|---|---|
| PP_0455 | 0.102632 | *rplD* | 50S ribosomal protein L4 |
| PP_0464 | 0.097197 | *rplN* | 50S ribosomal protein L14 |
| PP_0452 | 0.097141 | *tufB* | Elongation factor Tu-B |

Differentially active condition: growth phase, alcohol supplementation

#### 8. Zur iModulon (*n*=33, explained variance: 1.9%)

Category: Regulatory iModulon (MR and RR of 0.52 and 1.00, regulon discovery)

Inferred function: metal (zinc) homeostasis

Regulator: Zur (PP_0119)

Top five genes with highest weights:

| **Locus tag** | **Gene weight** | **Gene name** | **Gene product** |
|---|---|---|---|
| PP_5361 | 0.308422 | *PP_5361* | 47 kDa protein |
| PP_5360 | 0.266122 | *PP_5360* | conserved protein of unknown function |
| PP_0508 | 0.264463 | *PP_0508* | conserved exported protein of unknown function |
| PP_1446 | 0.257639 | *PP_1446* | TonB-dependent receptor |
| PP_3325 | 0.257514 | *PP_3325* | putative Outer membrane ferric siderophore receptor |

Differentially active condition: Zn ion, different sugars

#### 9. Uncharacterized-3 (*n*=11, explained variance: 1.8%)

Category: Uncharacterized iModulon

Top five genes with highest weights:

| Locus tag | Gene weight | Gene name | Gene product |
|---|---|---|---|
| PP_1661 | 0.178171 | *PP_1661* | putative Dehydrogenase subunit |
| PP_1660 | 0.176987 | *PP_1660* | conserved protein of unknown function |
| PP_1659 | 0.172653 | *PP_1659* | conserved exported protein of unknown function |
| PP_0806 | 0.141088 | *PP_0806* | putative Surface adhesion protein |
| PP_1245 | 0.140768 | *PP_1245* | conserved exported protein of unknown function |

Differentially active condition: M9 media adapted strains, glycolaldehyde addition

**10. FnrA-2 (*n*=18, explained variance: 1.7%)**

Category: Regulatory iModulon (MR and RR of 0.17 and 0.27, poorly-matched)

Inferred function: Metal (iron) homeostasis

Regulator: FnrA (PP_4265, also known as Anr)

Top five genes with highest weights:

| Locus tag | Gene weight | Gene name | Gene product |
|---|---|---|---|
| PP_1149 | 0.283627 | *PP_1149* | putative protein of unknown function |
| PP_5598 | 0.198975 | *PP_5598* | conserved protein of unknown function |
| PP_0504 | 0.1869 | *oprG* | Outer membrane protein OprG |
| PP_5392 | 0.17743 | *PP_5392* | conserved hypothetical protein containing WD40/YVTN repeat domain |
| PP_0273 | 0.166103 | *PP_0273* | conserved protein of unknown function |

Differentially active condition: M9 media adapted strains, glycolaldehyde addition

**11. PcaR iModulon (*n*=45, explained variance: 1.6%)**

Category: Regulatory iModulon (MR and RR of 0.20 and 1.00, regulon discovery)

Inferred function: aromatic acid catabolism

Regulator: PcaR (PP_1375), VanR (PP_3378), PP_3359

Top five genes with highest weights:

| **Locus tag** | **Gene weight** | **Gene name** | **Gene product** |
|---|---|---|---|
| PP_3357 | 0.198763 | *vdh* | vanillin dehydrogenase |
| PP_3350 | 0.197959 | *PP_3350* | conserved exported protein of unknown function |
| PP_3356 | 0.196945 | *fcs* | feruloyl-CoA-synthetase |
| PP_3358 | 0.193021 | *PP_3358* | hydroxycinnamoyl-CoA hydratase-lyase |
| PP_3355 | 0.192738 | *PP_3355* | Beta-ketothiolase |

Differentially active condition: M9 media adapted strains, glycolaldehyde addition

**12. Osmotic stress-2 iModulon (*n*=31, explained variance: 1.6%)**

Category: Functional iModulon

Inferred function: Osmotic stress mitigation

Regulator: Unknown

Top five genes with highest weights:

| **Locus tag** | **Gene weight** | **Gene name** | **Gene product** |
|---|---|---|---|
| PP_4707 | 0.154113 | *PP_4707* | OsmY-related protein |
| PP_3241 | 0.149571 | *PP_3241* | conserved exported protein of unknown function |

| | | | |
|---|---|---|---|
| PP_5592 | 0.147267 | *PP_5592* | conserved protein of unknown function |
| PP_1750 | 0.141363 | *asnB* | asparagine synthetase |
| PP_0869 | 0.140582 | *yehW* | osmoprotectant ABC transporter permease subunit |

Differentially active condition: NaCl addition

**13. InfA iModulon (*n*=4, explained variance: 1.5%)**

Category: Functional iModulon

Inferred function: Translation

Regulator: Unknown

Top four genes with highest weights:

| **Locus tag** | **Gene weight** | **Gene name** | **Gene product** |
|---|---|---|---|
| PP_4007 | 0.117582 | *infA* | Translation initiation factor IF-1 |
| PP_0686 | 0.086448 | *yjdM* | conserved protein of unknown function |
| PP_2220 | 0.08102 | *PP_2220* | C4-type zinc finger protein%2C DksA/TraR family |
| PP_5204 | 0.079879 | *PP_5204* | conserved protein of unknown function |

Differentially active condition: Long-term fermentation, oxidative stress

**14. Phage-2 iModulon (*n*=47, explained variance: 1.3%)**

Category: Functional iModulon

Inferred function: Phage related protein synthesis

Regulator: Unknown

Top five genes with highest weights:

| **Locus tag** | **Gene weight** | **Gene name** | **Gene product** |
|---|---|---|---|
| PP_1575 | 0.178634 | *PP_1575* | conserved protein of unknown function |
| PP_1574 | 0.171357 | *PP_1574* | conserved protein of unknown function |
| PP_1568 | 0.171082 | *PP_1568* | conserved protein of unknown function |
| PP_1572 | 0.168582 | *PP_1572* | conserved protein of unknown function |
| PP_1573 | 0.16823 | *PP_1573* | putative Major tail protein |

Differentially active condition: xylose minimal medium adaptation, aeration

#### 15. FleQ/AmrZ iModulon (*n*=44, explained variance: 1.3%)

Category: Regulatory iModulon (MR and RR of 0.84 and 0.12, regulon subset)

Inferred function: Flagella synthesis

Regulator: FleQ (PP_4373)

Top five genes with highest weights:

| **Locus tag** | **Gene weight** | **Gene name** | **Gene product** |
|---|---|---|---|
| PP_4370 | 0.17128 | *fliE* | Flagellar hook-basal body complex protein FliE |
| PP_4394 | 0.155183 | *flgA* | flagella basal body P-ring formation protein |
| PP_4359 | 0.14709 | *fliL* | Flagellar protein FliL |
| PP_4386 | 0.141075 | *flgF* | flagellar basal-body rod protein FlgF |

| | | | |
|---|---|---|---|
| PP_4358 | 0.140044 | *fliM* | flagellar synthesis%2C switching and energizing component |

Differentially active condition: *fleQ* deletion, *wspR* overexpression, osmotic stress

**16. FliA iModulon (*n*=29, explained variance: 1.2%)**

Category: Regulatory iModulon (MR and RR of 0.17 and 0.63, regulon discovery)

Inferred function: Cell mobility, flagella synthesis

Regulator: FliA (PP_4341)

Top five genes with highest weights:

| **Locus tag** | **Gene weight** | **Gene name** | **Gene product** |
|---|---|---|---|
| PP_4378 | 0.151963 | *fliC* | flagellin%2C filament structural protein |
| PP_4377 | 0.136108 | *PP_4377* | putative Flagellin FlaG |
| PP_1371 | 0.12591 | *pctA* | Methyl-accepting chemotaxis protein PctA |
| PP_5020 | 0.123337 | *PP_5020* | putative methyl-accepting chemotaxis protein |
| PP_1828 | 0.1227 | *PP_1828* | conserved protein of unknown function |

Differentially active condition: *fleQ* deletion, *wspR* overexpression, serine utilization

**17. PtxS iModulon (*n*=11, explained variance: 1.2%)**

Category: Regulatory iModulon (MR and RR of 0.73 and 1.00, well-matched)

Inferred function: Sugar catabolism

Regulator: PtxS (PP_3380), GnuR (PP_3415)

Top five genes with highest weights:

| Locus tag | Gene weight | Gene name | Gene product |
|---|---|---|---|
| PP_3378 | 0.334587 | *kguK* | putative 2-ketogluconokinase |
| PP_3377 | 0.331743 | *kguT* | 2-ketogluconate transporter%2C putative |
| PP_3379 | 0.327965 | *kguE* | putative epimerase |
| PP_3376 | 0.292836 | *ptxD* | putative phosphonate dehydrogenase |
| PP_3384 | 0.249982 | *PP_3384* | gluconate 2-dehydrogenase gamma subunit |

Differentially active condition: substrate type (non glucose condition)

**18. Null-1 iModulon (*n*=0)**

**19. Uncharacterized-2 iModulon (*n*=11, explained variance: 1.2%)**

Category: Uncharacterized iModulon

Inferred function: unknown

Regulator: unknown

Top five genes with highest weights:

| Locus tag | Gene weight | Gene name | Gene product |
|---|---|---|---|
| PP_0023 | 0.11579 | *PP_0023* | conserved hypothetical protein |
| PP_2322 | 0.105257 | *oprI* | Major outer membrane lipoprotein |
| PP_3704 | 0.104491 | *PP_3704* | conserved protein of unknown function |
| PP_0380 | 0.103003 | *pqqA* | Coenzyme PQQ synthesis protein A |

| | | | |
|---|---|---|---|
| PP_1249 | 0.102727 | *PP_1249* | conserved protein of unknown function%2C DUF4223 family |

Differentially active condition: alcohol addition, *mfsR* overexpression,

#### 20. TurA-2 iModulon (*n*=9, explained variance: 1.1%)

Category: Regulatory iModulon (MR and RR of 0.56 and 0.02, poorly-matched)

Inferred function: Chaperone

Regulator: TurA (PP_1366)

Top five genes with highest weights:

| **Locus tag** | **Gene weight** | **Gene name** | **Gene product** |
|---|---|---|---|
| PP_5000 | 0.169816 | *hslV* | peptidase component of the ATP-dependent HslVU protease |
| PP_5001 | 0.165801 | *hslU* | protease HslVU ATPase component |
| PP_4728 | 0.155785 | *grpE* | Protein GrpE |
| PP_1360 | 0.153978 | *groS* | 10 kDa chaperonin |
| PP_4179 | 0.149527 | *htpG* | Chaperone protein HtpG |

Differentially active condition: alcohol addition, osmotic stress

#### 21. CysB/PP_0204 iModulon (*n*=29, explained variance: 1.1%)

Category: Regulatory iModulon (MR and RR of 0.10 and 0.38, poorly-matched)

Inferred function: Sulfur metabolism

Regulator: CysB (PP_2327), PP_0204

Top five genes with highest weights:

| Locus tag | Gene weight | Gene name | Gene product |
|---|---|---|---|
| PP_5172 | 0.200838 | *PP_5172* | hypothetical protein |
| PP_5171 | 0.187942 | *sbp-II* | sulfate ABC transporter |
| PP_0233 | 0.167427 | *tauA* | taurine ABC transporter periplasmic binding subunit |
| PP_0236 | 0.160499 | *ssuE* | NAD(P)H-dependent FMN reductase subunit |
| PP_5166 | 0.15554 | *PP_5166* | Sigma-54 dependent transcriptional regulator |

Differentially active condition: glycolaldehyde addition, oxidative stress

#### 22. Osmotic stress-1 iModulon (*n*=29, explained variance: 1.0%)

Category: Functional iModulon

Inferred function: Sulfur metabolism

Regulator: unknown

Top five genes with highest weights:

| Locus tag | Gene weight | Gene name | Gene product |
|---|---|---|---|
| PP_3929 | 0.158821 | *PP_3929* | conserved protein of unknown function |
| PP_3631 | 0.136082 | *htrG* | putative signal transduction protein |
| PP_5038 | 0.130099 | *PP_5038* | conserved lipoprotein of unknown function |
| PP_0837 | 0.125109 | *ycfJ* | putative regulator of flagellation |
| PP_5702 | 0.122871 | *PP_5702* | conserved protein of unknown function |

Differentially active condition: isopentanol or NaCl addition, adapted strains in the presence of an ionic liquid

#### 23. Uncharacterized-6 iModulon (*n*=80, explained variance: 1.0%)

Category: Uncharacterized iModulon

Inferred function: unknown

Regulator: unknown

Top five genes with highest weights:

| Locus tag | Gene weight | Gene name | Gene product |
|---|---|---|---|
| PP_5610 | 0.11895 | *PP_5610* | protein of unknown function |
| PP_4447 | 0.114413 | *PP_4447* | conserved protein of unknown function |
| PP_0640 | 0.112628 | *PP_0640* | conserved protein of unknown function |
| PP_1959 | 0.112607 | *PP_1959* | conserved protein of unknown function |
| PP_5491 | 0.111978 | *PP_5491* | conserved protein of unknown function with SEC-C motif domain |

Differentially active condition: *mfsR* overexpression

#### 24. Null-2 iModulon

#### 25. PplR1 iModulon (*n*=22, explained variance: 0.94%)

Category: Regulatory iModulon (MR and RR of 0.77 and 1.00, well-matched)

Inferred function: Light inducible iModulon

Regulator: PplR1 (PP_0740)

Top five genes with highest weights:

| Locus tag | Gene weight | Gene name | Gene product |
|---|---|---|---|
| PP_2730 | 0.245689 | *PP_2730* | putative Lipoprotein |

| | | | |
|---|---|---|---|
| PP_2729 | 0.237205 | *PP_2729* | conserved protein of unknown function |
| PP_2731 | 0.229905 | *PP_2731* | conserved protein of unknown function |
| PP_2736 | 0.229815 | *PP_2736* | conserved protein of unknown function |
| PP_2737 | 0.220654 | *PP_2737* | Oxidoreductase%2C short-chain dehydrogenase/reductase family |

Differentially active condition: sugar type

#### 26. Uncharacterized-1 iModulon (*n*=6, explained variance: 0.94%)

Category: Regulatory iModulon

Inferred function: unknown

Regulator: unknown

Top five genes with highest weights:

| **Locus tag** | **Gene weight** | **Gene name** | **Gene product** |
|---|---|---|---|
| PP_2857 | 0.376915 | *PP_2857* | conserved exported protein of unknown function |
| PP_2856 | 0.32534 | *PP_2856* | conserved exported protein of unknown function |
| PP_2858 | 0.313172 | *PP_2858* | conserved exported protein of unknown function |
| PP_2855 | 0.22664 | *PP_2855* | conserved exported protein of unknown function |
| PP_2854 | 0.2035 | *PP_2854* | conserved exported protein of unknown function |

Differentially active condition: substrate type (aromatic compounds, sugars)

#### 27. GltR-II iModulon (*n*=7, explained variance: 0.90%)

Category: Regulatory iModulon (MR and RR of 0.71 and 0.63, well-matched)

Inferred function: glucose metabolism

Regulator: GltR-II (PP_1012)

Top five genes with highest weights:

| **Locus tag** | **Gene weight** | **Gene name** | **Gene product** |
|---|---|---|---|
| PP_1015 | 0.390153 | *gtsA* | mannose/glucose ABC transporter%2C glucose-binding periplasmic protein |
| PP_1018 | 0.372419 | *gtsD* | mannose/glucose ABC transporter - ATP binding subunit |
| PP_1017 | 0.368212 | *gtsC* | mannose/glucose ABC transporter%2C permease protein |
| PP_1016 | 0.356304 | *gtsB* | mannose/glucose ABC transporter%2C permease protein |
| PP_1019 | 0.333482 | *oprB-I* | carbohydrate-selective porin |

Differentially active condition: substrate type (aromatic compounds, sugars)

#### 28. Pentose and glucuronate iModulon (*n*=7, explained variance: 0.86%)

Category: Functional iModulon

Inferred function: glucose metabolism

Regulator: unknown

Top five genes with highest weights:

| **Locus tag** | **Gene weight** | **Gene name** | **Gene product** |
|---|---|---|---|
| PP_2585 | 0.26249 | *PP_2585* | Alpha-ketoglutaric semialdehyde dehydrogenase |
| PP_2834 | 0.260131 | *PP_2834* | Putative D-galactarate dehydratase/Altronate dehydratase |

| | | | |
|---|---|---|---|
| PP_2835 | 0.245469 | *PP_2835* | putative oxidoreductase |
| PP_2836 | 0.239918 | *PP_2836* | putative 2-keto-3-deoxyxylonate dehydratase |
| PP_2837 | 0.232254 | *PP_2837* | putative uronate transporter |

Differentially active condition: xylose utilization

#### 29. Uncharacterized-7 iModulon (*n*=7, explained variance: 0.86%)

Category: Functional iModulon

Inferred function: unknown

Regulator: unknown

Top five genes with highest weights:

| Locus tag | Gene weight | Gene name | Gene product |
|---|---|---|---|
| PP_5421 | 0.128318 | *PP_5421* | conserved protein of unknown function |
| PP_0014 | 0.122861 | *PP_0014* | putative transposase |
| PP_4462 | 0.117422 | *PP_4462* | putative 4-hydroxy-4-methyl-2-oxoglutarate aldolase |
| PP_4465 | 0.116105 | *PP_4465* | putative Porin |
| PP_4464 | 0.114284 | *PP_4464* | Transcriptional regulator%2C LysR family |

Differentially active condition: NaCl addition, adaptation for galactose utilization

#### 30. Exporters (*n*=7, explained variance: 0.81%)

Category: Functional iModulon

Inferred function: multidrug efflux

Regulator: unknown

Top five genes with highest weights:

| Locus tag | Gene weight | Gene name | Gene product |
|---|---|---|---|
| PP_3425 | 0.311766 | *PP_3425* | Efflux transporter RND family%2C MFP subunit |
| PP_3426 | 0.291577 | *mexF* | Multidrug efflux RND transporter MexF |
| PP_3427 | 0.286734 | *oprN* | Multidrug efflux RND outer membrane protein OprN |
| PP_4858 | 0.280781 | *PP_4858* | conserved exported protein of unknown function |
| PP_5496 | 0.26293 | *PP_5496* | conserved exported protein of unknown function |

Differentially active condition: deletion of *fleQ*, overexpression of *wspR*, altered *turA* and *turB* expression

**31. LiuR iModulon (*n*=12, explained variance: 0.81%)**

Category: Regulatory iModulon (MR and RR of 0.33 and 0.57, poorly-matched)

Inferred function: Ribose and lipid catabolism

Regulator: LiuR (PP_3539)

Top five genes with highest weights:

| Locus tag | Gene weight | Gene name | Gene product |
|---|---|---|---|
| PP_5313 | 0.10705 | *hupA* | DNA-binding protein HU-alpha |
| PP_2455 | 0.104793 | *rbsA-I* | ribose ABC transporter - ATP-binding subunit |
| PP_4401 | 0.095879 | *bkdAA* | branched-chain alpha-keto acid dehydrogenase complex%2C alpha subunit |

| | | | |
|---|---|---|---|
| PP_3726 | 0.095072 | *PP_3726* | Enoyl-CoA hydratase/isomerase family protein |
| PP_2454 | 0.092143 | *rbsB* | ribose ABC transporter%2C periplasmic ribose-binding subunit |

Differentially active condition: growth phase, butanol addition

#### 32. GbdR iModulon (*n*=12, explained variance: 0.81%)

Category: Regulatory iModulon (MR and RR of 0.47 and 0.55, poorly-matched)

Inferred function: Glycine Betaine Catabolism

Regulator: GbdR (PP_0298)

Top five genes with highest weights:

| Locus tag | Gene weight | Gene name | Gene product |
|---|---|---|---|
| PP_0310 | 0.13648 | *dgcA* | putative dimethylglycine dehydrogenase subunit |
| PP_0309 | 0.133286 | *PP_0309* | conserved protein of unknown function |
| PP_0328 | 0.121385 | *fdhA* | formaldehyde dehydrogenase |
| PP_0315 | 0.119117 | *gbcA* | putative glycine-betaine dioxygenase subunit |
| PP_4619 | 0.117153 | *hmgC* | Maleylacetoacetate isomerase |

Differentially active condition: growth phase

#### 33. TCA cycle iModulon (*n*=12, explained variance: 0.81%)

Category: Functional iModulon

Inferred function: TCA cycle

Regulator: unknown

Top five genes with highest weights:

| Locus tag | Gene weight | Gene name | Gene product |
|---|---|---|---|
| PP_0354 | 0.196179 | *PP_0354* | CBS domain protein |
| PP_0353 | 0.19198 | *PP_0353* | Exonuclease |
| PP_3179 | 0.168713 | *PP_3179* | Transcriptional regulator%2C LysR family |
| PP_3014 | 0.159936 | *PP_3014* | conserved exported protein of unknown function |
| PP_1742 | 0.15088 | *yjcH* | conserved inner membrane protein of unknown function |

Differentially active condition: sugar types, oxidative stress, alcohol addition

**34. FleQ/Fur iModulon (*n*=36, explained variance: 0.76%)**

Category: Regulatory iModulon (MR and RR of 0.72 and 0.09, regulon subset)

Inferred function: metal (iron) homeostasis

Regulator: FleQ (PP_4373), Fur (PP_4730)

Top five genes with highest weights:

| Locus tag | Gene weight | Gene name | Gene product |
|---|---|---|---|
| PP_1083 | 0.169198 | *PP_1083* | (2Fe-2S)-binding protein |
| PP_4070 | 0.167086 | *PP_4070* | conserved protein of unknown function |
| PP_0352 | 0.160829 | *PP_0352* | RNA polymerase sigma-70 factor%2C ECF subfamily |
| PP_5306 | 0.152129 | *exbB* | Biopolymer transport protein ExbB |
| PP_5307 | 0.145702 | *exbD* | TonB-gated outer membrane transporter - gating inner membrane protein |

Differentially active condition: minimal media adaptation

#### 35. GlnG iModulon (*n*=11, explained variance: 0.75%)

Category: Regulatory iModulon (MR and RR of 0.82 and 0.12, regulon subset)

Inferred function: Amino acid transportation

Regulator: GlnG (PP_5048)

Top five genes with highest weights:

| Locus tag | Gene weight | Gene name | Gene product |
|---|---|---|---|
| PP_1297 | 0.220819 | *yhdW* | putative amino-acid ABC transporter-binding protein YhdW |
| PP_1298 | 0.180647 | *yhdX* | putative amino acid ABC transporter - permease subunit |
| PP_4867 | 0.175711 | *PP_4867* | branched-chain amino acid ABC transporter%2C periplasmic amino acid-binding protein (braC-like) |
| PP_1299 | 0.175517 | *yhdY* | putative amino acid ABC transporter - membrane subunit |
| PP_4864 | 0.159589 | *braF* | High-affinity branched-chain amino acid transport ATP-binding protein BraF |

Differentially active condition: deletion of *crcZ* and *crcY*, utilization of galactose, xylose, or serine

#### 36. GclR iModulon (*n*=14, explained variance: 0.73%)

Category: Regulatory iModulon (MR and RR of 0.71 and 0.53, regulon subset)

Inferred function: C2 carbon metabolism

Regulator: GclR (PP_5048)

Top five genes with highest weights:

| Locus tag | Gene weight | Gene name | Gene product |
|---|---|---|---|

| | | | |
|---|---|---|---|
| PP_4288 | 0.147197 | *allA* | ureidoglycolate lyase |
| PP_4286 | 0.14113 | *puuE* | allantoinase |
| PP_0596 | 0.138668 | *PP_0596* | Omega-amino acid--pyruvate aminotransferase |
| PP_4287 | 0.13397 | *pucL* | 2-oxo-4-hydroxy-4-carboxy-5-ureidoimidazoline decarboxylase |
| PP_0597 | 0.113831 | *mmsA-I* | methylmalonate-semialdehyde dehydrogenase |

Differentially active condition: long term cultivation, isopentanol addition

#### 37. PP_2260-3 iModulon (*n*=5, explained variance: 0.73%)

Category: Functional iModulon

Inferred function: PP_2260-3, sugar transport

Regulator: unknown

Top five genes with highest weights:

| **Locus tag** | **Gene weight** | **Gene name** | **Gene product** |
|---|---|---|---|
| PP_2261 | 0.245515 | *PP_2261* | Sugar ABC transporter ATP-binding protein |
| PP_2260 | 0.24189 | *PP_2260* | putative glycerol-phosphate ABC transporter ATP-binding protein |
| PP_5505 | 0.21717 | *PP_5505* | conserved transmembrane protein of unknown function |
| PP_2262 | 0.217027 | *PP_2262* | sugar ABC transporter permease protein |
| PP_2263 | 0.149185 | *PP_2263* | Sugar ABC transporter permease protein |

Differentially active condition: substrate type, growth phase

#### 38. GcsR iModulon (*n*=4, explained variance: 0.73%)

Category: Regulatory iModulon (MR and RR of 1.00 and 1.00, well-matched)

Inferred function: Glycine metabolism

Regulator: GcsR (PP_0997)

Four genes with their weights:

| **Locus tag** | **Gene weight** | **Gene name** | **Gene product** |
|---|---|---|---|
| PP_0989 | 0.250466 | *gcvH-I* | glycine cleavage system H protein 1 |
| PP_0988 | 0.148719 | *gcvP-I* | glycine dehydrogenase |
| PP_0987 | 0.126096 | *tdcG-II* | L-serine dehydratase |
| PP_0986 | 0.119095 | *gcvT-I* | aminomethyltransferase |

Differentially active condition: aromatic acid utilization, growth phase

#### 39. OxyR iModulon (*n*=4, explained variance: 0.72%)

Category: Regulatory iModulon (MR and RR of 0.25 and 0.75, regulon discovery)

Inferred function: Oxidative stress

Regulator: OxyR (PP_5309)

Top five genes with highest weights:

| **Locus tag** | **Gene weight** | **Gene name** | **Gene product** |
|---|---|---|---|
| PP_0481 | 0.16214 | *katA* | Catalase |
| PP_2440 | 0.150477 | *ahpF* | Alkyl hydroperoxide reductase subunit F |
| PP_1834 | 0.146094 | *PP_1834* | conserved protein of unknown function |

| | | | |
|---|---|---|---|
| PP_1836 | 0.114947 | *PP_1836* | putative metal transporter%2C ZIP family |
| PP_2309 | 0.112158 | *PP_2309* | conserved protein of unknown function |

Differentially active condition: $H_2O_2$ or isopentanol addition

**40. FnrA-1 iModulon (*n*=16, explained variance: 0.71%)**

Category: Regulatory iModulon (MR and RR of 0.19 and 0.27, poorly-matched)

Inferred function: Redox state homeostasis

Regulator: FnrA (PP_4265)

Top five genes with highest weights:

| Locus tag | Gene weight | Gene name | Gene product |
|---|---|---|---|
| PP_4253 | 0.176765 | *ccoP-I* | cytochrome c oxidase subunit cbb3-type |
| PP_4252 | 0.170987 | *ccoQ-I* | cytochrome c oxidase subunit cbb3-type |
| PP_4251 | 0.16415 | *ccoO-I* | cytochrome c oxidase subunit cbb3-type |
| PP_1000 | 0.15475 | *arcB* | ornithine carbamoyltransferase catabolic |
| PP_5734 | 0.136101 | *PP_5734* | conserved hypothetical protein |

Differentially active condition: $H_2O_2$ or isopentanol addition,

**41. Acetate iModulon (*n*=10, explained variance: 0.67%)**

Category: Functional iModulon

Inferred function: Acetate stress response

Regulator: unknown

Top five genes with highest weights:

| Locus tag | Gene weight | Gene name | Gene product |
|---|---|---|---|
| PP_4488 | 0.354262 | *PP_4488* | conserved exported protein of unknown function |
| PP_0372 | 0.275029 | *aruC* | Acetylornithine aminotransferase 2 |
| PP_0269 | 0.203705 | *PP_0269* | putative Glutamate synthase large subunit |
| PP_5458 | 0.202431 | *PP_5458* | conserved exported protein of unknown function |
| PP_1265 | 0.200728 | *PP_1265* | conserved protein of unknown function |

Differentially active condition: acetate addition, zinc ion

#### 42. PvdS iModulon (*n*=36, explained variance: 0.67%)

Category: Regulatory iModulon (MR and RR of 0.33 and 0.92, regulon discovery)

Inferred function: Pyoverdine biosynthesis

Regulator: PvdS (PP_4244)

Top five genes with highest weights:

| Locus tag | Gene weight | Gene name | Gene product |
|---|---|---|---|
| PP_3796 | 0.182534 | *pvdA* | L-ornithine 5-monooxygenase |
| PP_4217 | 0.171365 | *fpvA* | outer membrane ferripyoverdine receptor FpvA%2C TonB-dependent |
| PP_4223 | 0.162897 | *pvdH* | diaminobutyrate-2-oxoglutarate transaminase |
| PP_3808 | 0.162794 | *PP_3808* | Antibiotic synthesis protein MbtH |
| PP_3807 | 0.158096 | *PP_3807* | Thioesterase |

Differentially active condition: *turA* deletion, lignin hydrolysate addition, aeration, acetate addition

#### 43. HexR iModulon (*n*=9, explained variance: 0.64%)

Category: Regulatory iModulon (MR and RR of 1.00 and 1.00, well-matched)

Inferred function: Sugar catabolism

Regulator: HexR (PP_1021)

Top five genes with highest weights:

| Locus tag | Gene weight | Gene name | Gene product |
|---|---|---|---|
| PP_1022 | 0.280974 | *zwfA* | glucose 6-phosphate-1-dehydrogenase |
| PP_1023 | 0.276182 | *pgl* | 6-phosphogluconolactonase |
| PP_1009 | 0.256776 | *gapA* | glyceraldehyde-3-phosphate dehydrogenase |
| PP_1024 | 0.255461 | *eda* | KHG/KDPG aldolase |
| PP_1010 | 0.194919 | *edd* | phosphogluconate dehydratase |

Differentially active condition: hexose condition

#### 44. T6SS iModulon (*n*=15, explained variance: 0.64%)

Category: Functional iModulon

Inferred function: T6SS system

Regulator: unknown

Top five genes with highest weights:

| Locus tag | Gene weight | Gene name | Gene product |
|---|---|---|---|
| PP_3092 | 0.192605 | *PP_3092* | conserved protein of unknown function |

| | | | |
|---|---|---|---|
| PP_3089 | 0.190031 | *PP_3089* | conserved protein of unknown function |
| PP_3100 | 0.186974 | *PP_3100* | conserved protein of unknown function |
| PP_3093 | 0.178761 | *PP_3093* | conserved protein of unknown function |
| PP_3094 | 0.178246 | *PP_3094* | conserved exported protein of unknown function |

Differentially active condition: growth phase, NaCl addition, minimal media adaptation

#### 45. Redox iModulon (*n*=3, explained variance: 0.63%)

Category: Functional iModulon

Inferred function: Redox state homeostasis

Regulator: unknown

Three genes with their weights:

| Locus tag | Gene weight | Gene name | Gene product |
|---|---|---|---|
| PP_3332 | 0.426734 | *PP_3332* | putative cytochrome c-type protein |
| PP_3333 | 0.339726 | *PP_3333* | conserved protein of unknown function |
| PP_5743 | 0.214019 | *PP_5743* | putative TonB-dependent receptor protein |

Differentially active condition: substrate type

#### 46. Multiple stress-2 iModulon (*n*=3, explained variance: 0.63%)

Category: Functional iModulon

Inferred function: Redox state homeostasis

Regulator: unknown

Three genes with their weights:

| Locus tag | Gene weight | Gene name | Gene product |
|---|---|---|---|
| PP_3232 | 0.195903 | *PP_3232* | Acetyltransferase%2C GNAT family |
| PP_3236 | 0.163384 | *PP_3236* | putative Lipoprotein OprI |
| PP_3289 | 0.149117 | *PP_3289* | Acetyltransferase%2C GNAT family |
| PP_2745 | 0.136787 | *PP_2745* | Universal stress protein family |
| PP_3235 | 0.125323 | *PP_3235* | conserved protein of unknown function |

Differentially active condition: substrate type

#### 47. Uncharacterized-9 iModulon (*n*=2, explained variance: 0.61%)

Category: Uncharacterized iModulon

Inferred function: unknown

Regulator: unknown

Top five genes with highest weights:

| Locus tag | Gene weight | Gene name | Gene product |
|---|---|---|---|
| PP_0153 | 0.12939 | *PP_0153* | hypothetical protein |
| PP_4615 | 0.118746 | *PP_4615* | conserved membrane protein of unknown function |

Differentially active condition: substrate type

#### 48. Phage-1 iModulon (*n*=29, explained variance: 0.57%)

Category: Functional iModulon

Inferred function: Phage related

Regulator: unknown

Top five genes with highest weights:

| Locus tag | Gene weight | Gene name | Gene product |
|---|---|---|---|
| PP_3877 | 0.198433 | *PP_3877* | conserved protein of unknown function |
| PP_3870 | 0.159007 | *PP_3870* | Phage FluMu protein gp38 |
| PP_3876 | 0.156867 | *PP_3876* | conserved protein of unknown function |
| PP_3875 | 0.151366 | *PP_3875* | conserved protein of unknown function |
| PP_3871 | 0.144202 | *PP_3871* | conserved protein of unknown function |

Differentially active condition: *relA* deletion, xylose minimal medium adaptation

**49. PP_2034-7 iModulon (*n*=4, explained variance: 0.56%)**

Category: Functional iModulon

Inferred function: Benzoate catabolism

Regulator: unknown

Top five genes with highest weights:

| Locus tag | Gene weight | Gene name | Gene product |
|---|---|---|---|
| PP_2037 | 0.334823 | *PP_2037* | putative Aldolase |
| PP_2036 | 0.299485 | *PP_2036* | putative 4-hydroxy-tetrahydrodipicolinate synthase |

| | | | |
|---|---|---|---|
| PP_2035 | 0.232224 | *benE-I* | benzoate transport protein |
| PP_2034 | 0.190576 | *PP_2034* | conserved membrane protein of unknown function |

Differentially active condition: NaCl addition, lignin hydrolysate addition

**50. FruR iModulon (*n*=4, explained variance: 0.51%)**

Category: Regulatory iModulon (MR and RR of 0.75 and 0.60, regulon subset)

Inferred function: Fructose catabolism

Regulator: FruR (PP_0792, also known as Cra)

Top five genes with highest weights:

| Locus tag | Gene weight | Gene name | Gene product |
|---|---|---|---|
| PP_0794 | 0.251425 | *fruK* | 1-phosphofructokinase monomer |
| PP_0795 | 0.250917 | *fruA* | fructose PTS permease - IIBC component |
| PP_0793 | 0.225521 | *fruB* | Phosphotransferase system fructose-specific EI/HPr/EIIA components |
| PP_3443 | 0.150902 | *PP_3443* | putative glyceraldehyde-3-phosphate dehydrogenase |

Differentially active condition: hexose condition

**51. PsrA iModulon (*n*=13, explained variance: 0.51%)**

Category: Regulatory iModulon (MR and RR of 0.31 and 0.29, poorly-matched)

Inferred function: Fatty acid catabolism

Regulator: PsrA (PP_2144)

Top five genes with highest weights:

| Locus tag | Gene weight | Gene name | Gene product |
|---|---|---|---|
| PP_3755 | 0.156886 | *hbd* | 3-hydroxybutyryl-CoA dehydrogenase |
| PP_3756 | 0.14779 | *PP_3756* | TetR family transcriptional regulator |
| PP_3754 | 0.146215 | *bktB* | Beta-ketothiolase BktB |
| PP_5008 | 0.142388 | *PP_5008* | Poly granule-associated protein |
| PP_2136 | 0.135154 | *fadB* | enoyl-CoA hydratase/3-hydroxyacyl-CoA dehydrogenase |

Differentially active condition: myristic acid utilization

#### 52. AmrZ iModulon (*n*=12, explained variance: 0.48%)

Category: Regulatory iModulon (MR and RR of 1.00 and 0.09, regulon subset)

Inferred function: Alginate biosynthesis

Regulator: AmrZ (PP_4470)

Top five genes with highest weights:

| Locus tag | Gene weight | Gene name | Gene product |
|---|---|---|---|
| PP_1279 | 0.246248 | *algJ* | Probable alginate O-acetylase AlgJ |
| PP_1278 | 0.243565 | *algF* | Alginate biosynthesis protein AlgF |
| PP_1288 | 0.243184 | *algD* | GDP-mannose 6-dehydrogenase |
| PP_1286 | 0.204913 | *alg44* | Alginate biosynthesis protein Alg44 |
| PP_1282 | 0.195357 | *algX* | Alginate biosynthesis protein AlgX |

Differentially active condition: substrate type, alcohol addition, NaCl addition, minimal medium adaptation

#### 53. BenR iModulon (*n*=10, explained variance: 0.48%)

Category: Regulatory iModulon (MR and RR of 0.40 and 1.00, regulon discovery)

Inferred function: Alginate biosynthesis

Regulator: AmrZ (PP_4470)

Top five genes with highest weights:

| **Locus tag** | **Gene weight** | **Gene name** | **Gene product** |
|---|---|---|---|
| PP_3162 | 0.2592 | *benB* | benzoate 1,2-dioxygenase subunit beta |
| PP_3163 | 0.242219 | *benC* | benzoate 1,2-dioxygenase electron transfer component |
| PP_3166 | 0.230763 | *catA-II* | catechol 1,2-dioxygenase |
| PP_3164 | 0.228605 | *benD* | 1,6-dihydroxycyclohexa-2,4-diene-1-carboxylate dehydrogenase |
| PP_3161 | 0.22578 | *benA* | benzoate 1,2-dioxygenase subunit alpha |

Differentially active condition: substrate type, *turA* deletion, alcohol addition

#### 54. Acetate iModulon (*n*=4, explained variance: 0.47%)

Category: Functional iModulon

Inferred function: Acetate utilization

Regulator: unknown

Top five genes with highest weights:

| **Locus tag** | **Gene weight** | **Gene name** | **Gene product** |
|---|---|---|---|
| PP_1742 | 0.309203 | *yjcH* | conserved inner membrane protein of unknown function |

| | | | |
|---|---|---|---|
| PP_0354 | 0.264953 | *PP_0354* | CBS domain protein |
| PP_1743 | 0.243364 | *actP-I* | acetate permease |
| PP_4487 | 0.171203 | *acsA-I* | acetyl-CoA synthetase |

Differentially active condition: alcohol addition, substrate type, $H_2O_2$ addition, zinc ion addition

#### 55. AcoR iModulon (*n*=5, explained variance: 0.46%)

Category: Regulatory iModulon (MR and RR of 1.00 and 0.71, well-matched)

Inferred function: Acetoin catabolism

Regulator: AcoR (PP_0557)

Top five genes with highest weights:

| Locus tag | Gene weight | Gene name | Gene product |
|---|---|---|---|
| PP_0555 | 0.314694 | *acoA* | Acetoin:2,6-dichlorophenolindophenol oxidoreductase subunit alpha |
| PP_0553 | 0.264192 | *acoC* | Dihydrolipoyllysine-residue acetyltransferase component of acetoin cleaving system |
| PP_0556 | 0.259332 | *PP_0556* | Acetoin catabolism protein |
| PP_0554 | 0.246452 | *acoB* | Acetoin:2,6-dichlorophenolindophenol oxidoreductase subunit beta |
| PP_0552 | 0.169372 | *bdhA* | 2,3-butanediol dehydrogenase |

Differentially active condition: *crc* deletion, substrate type

#### 56. Putrescine iModulon (*n*=7, explained variance: 0.42%)

Category: Functional iModulon

Inferred function: beta-Alanine metabolism

Putative regulator: PuuR (PP_5268)

Top five genes with highest weights:

| Locus tag | Gene weight | Gene name | Gene product |
|---|---|---|---|
| PP_3598 | 0.171721 | *PP_3598* | Peptidase C26 |
| PP_0596 | 0.148885 | *PP_0596* | Omega-amino acid--pyruvate aminotransferase |
| PP_2448 | 0.14226 | *puuB* | gamma-glutamylputrescine oxidase |
| PP_1229 | 0.132848 | *puuP* | putrescine permease |
| PP_0597 | 0.12652 | *mmsA-I* | methylmalonate-semialdehyde dehydrogenase |

Differentially active condition: growth phase, NaCl or $H_2O_2$ addition, *relA* deletion

**57. HutC iModulon (*n*=9, explained variance: 0.41%)**

Category: Regulatory iModulon (MR and RR of 0.67 and 0.75, well-matched)

Inferred function: beta-Alanine metabolism

Regulator: HutC (PP_5035)

Top five genes with highest weights:

| Locus tag | Gene weight | Gene name | Gene product |
|---|---|---|---|
| PP_5033 | 0.193536 | *hutU* | Urocanate hydratase |
| PP_5029 | 0.18606 | *hutG* | N-formylglutamate deformylase |
| PP_5036 | 0.18365 | *hutF* | probable formiminoglutamate deiminase |
| PP_5030 | 0.169608 | *hutI* | Imidazolonepropionase |

| | | | |
|---|---|---|---|
| PP_5032 | 0.166828 | *hutH* | Histidine ammonia-lyase |

Differentially active condition: growth phase, *finR* deletion, *wspR* overexpression

**58. PP_5350 iModulon (*n*=4, explained variance: 0.40%)**

Category: Regulatory iModulon (MR and RR of 0.75 and 0.43, regulon subset)

Inferred function: Glyoxylate and dicarboxylate metabolism

Regulator: PP_5350

Four genes with their weights:

| **Locus tag** | **Gene weight** | **Gene name** | **Gene product** |
|---|---|---|---|
| PP_2875 | 0.299032 | *PP_2875* | conserved protein of unknown function |
| PP_4116 | 0.295064 | *aceA* | isocitrate lyase |
| PP_5659 | 0.292753 | *PP_5659* | protein of unknown function |
| PP_0356 | 0.149902 | *glcB* | malate synthase G |

Differentially active condition: myristic acid utilization, acetate addition

**59. HmgR iModulon (*n*=4, explained variance: 0.40%)**

Category: Regulatory iModulon (MR and RR of 0.50 and 0.50, poorly-matched)

Inferred function: Tyrosine metabolism

Regulator: HmgR (PP_4622)

Four genes with their weights:

| **Locus tag** | **Gene weight** | **Gene name** | **Gene product** |
|---|---|---|---|

| | | | |
|---|---|---|---|
| PP_4621 | 0.243717 | *hmgA* | Homogentisate 1%2C2-dioxygenase |
| PP_3434 | 0.216344 | *PP_3434* | conserved protein of unknown function |
| PP_3433 | 0.182865 | *hpd* | 4-hydroxyphenylpyruvate dioxygenase |
| PP_4619 | 0.17266 | *hmgC* | Maleylacetoacetate isomerase |

Differentially active condition: butanol addition, substrate type

#### 60. MetR-I iModulon (*n*=6, explained variance: 0.38%)

Category: Regulatory iModulon (MR and RR of 0.83 and 0.50, regulon subset)

Inferred function: Homocysteine metabolism

Regulator: MetR-I (PP_1063)

Top five genes with highest weights:

| Locus tag | Gene weight | Gene name | Gene product |
|---|---|---|---|
| PP_2699 | 0.366821 | *PP_2699* | conserved protein of unknown function |
| PP_2698 | 0.331529 | *metE* | 5-methyltetrahydropteroyltriglutamate-homocysteine methyltransferase |
| PP_2697 | 0.29631 | *PP_2697* | putative oxidoreductase containing a flavin reductase domain |
| PP_4637 | 0.27101 | *PP_4637* | 5-methyltetrahydropteroyltriglutamate-homocysteine S-methyltransferase family protein |
| PP_2696 | 0.23556 | *metR-II* | DNA-binding transcriptional regulator homocysteine-binding |

Differentially active condition: substrate type

#### 61. PP_0615-8 iModulon (*n*=5, explained variance: 0.38%)

Category: Functional iModulon

Inferred function: branched amino acid transportation

Regulator: unknown

Top five genes with highest weights:

| Locus tag | Gene weight | Gene name | Gene product |
|---|---|---|---|
| PP_0618 | 0.198407 | *PP_0618* | Branched-chain amino acid ABC transporter%2C permease protein |
| PP_0616 | 0.195879 | *PP_0616* | Branched-chain amino acid ABC transporter%2C ATP binding protein |
| PP_0617 | 0.187816 | *PP_0617* | Branched-chain amino acid ABC transporter%2C permease protein |
| PP_0615 | 0.173664 | *PP_0615* | Branched-chain amino acid ABC transporter%2C ATP-binding protein |
| PP_0619 | 0.146524 | *PP_0619* | Branched-chain amino acid ABC transporter%2C periplasmic amino acid-binding protein |

Differentially active condition: substrate type

#### 62. Uncharacterized-8 iModulon (*n*=5, explained variance: 0.38%)

Category: Uncharacterized iModulon

Inferred function: unknown

Regulator: unknown

Top five genes with highest weights:

| Locus tag | Gene weight | Gene name | Gene product |
|---|---|---|---|
| PP_3336 | 0.330292 | *PP_3336* | conserved protein of unknown function |
| PP_4837 | 0.313869 | *PP_4837* | conserved protein of unknown function |

| | | | |
|---|---|---|---|
| PP_3335 | 0.256822 | *PP_3335* | conserved protein of unknown function |
| PP_4838 | 0.255625 | *oprC* | Outer membrane copper receptor OprC |
| PP_3330 | 0.219612 | *PP_3330* | putative Outer membrane ferric siderophore receptor |

Differentially active condition: substrate type

#### 63. ColR iModulon (*n*=5, explained variance: 0.38%)

Category: Regulatory iModulon (MR and RR of 0.0.27 and 0.27, poorly-matched)

Inferred function: Metal related response

Regulator: ColR (PP_0901)

Top five genes with highest weights:

| **Locus tag** | **Gene weight** | **Gene name** | **Gene product** |
|---|---|---|---|
| PP_0034 | 0.154268 | *PP_0034* | putative bactoprenol glycosyl-transferase from phage origin |
| PP_0036 | 0.147394 | *PP_0036* | putative transcriptional regulator |
| PP_0035 | 0.136522 | *PP_0035* | putative bactoprenol-linked glycose transferase |
| PP_0042 | 0.125864 | *PP_0042* | conserved exported protein of unknown function |
| PP_5138 | 0.116755 | *PP_5138* | putative transcriptional regulator LysR family |

Differentially active condition: zinc ion addition

#### 64. LldR iModulon (*n*=4, explained variance: 0.33%)

Category: Regulatory iModulon (MR and RR of 0.75 and 0.75, well-matched)

Inferred function: Lactate catabolism

Regulator: LldR (PP_4734)

Four genes with their weights:

| Locus tag | Gene weight | Gene name | Gene product |
|---|---|---|---|
| PP_4735 | 0.429437 | *lldP* | L-lactate permease |
| PP_4736 | 0.330067 | *lldD* | L-lactate dehydrogenase |
| PP_4737 | 0.287964 | *dld2* | D-lactate dehydrogenase |
| PP_1188 | 0.277198 | *dctA-I* | C4-dicarboxylate transport protein |

Differentially active condition: imipenem addition, citrate utilization

#### 65. Noise-5 iModulon

#### 66. Noise-6 iModulon

#### 67. AA transport iModulon (*n*=11, explained variance: 0.31%)

Category: Functional iModulon

Inferred function: Amino acid transportation

Regulator: unknown

Top five genes with highest weights:

| Locus tag | Gene weight | Gene name | Gene product |
|---|---|---|---|
| PP_3593 | 0.173571 | *PP_3593* | Amino acid ABC transporter periplasmic binding protein |
| PP_3596 | 0.172196 | *amaD* | D-lysine oxidase |
| PP_0883 | 0.161313 | *opdP* | glycine-glutamate dipeptide porin |

| | | | |
|---|---|---|---|
| PP_3594 | 0.157995 | *PP_3594* | Amino acid ABC transporter membrane protein |
| PP_3595 | 0.134629 | *PP_3595* | Amino acid ABC transporter membrane protein |

Differentially active condition: growth phase

**68. Genomic-2 iModulon (*n*=4, explained variance: 0.31%)**

Category: Genomic iModulon

Active condition: HNS study (deletion)

**69. Uncharacterized-5 iModulon (*n*=12, explained variance: 0.2872%)**

Category: Uncharacterized iModulon

Inferred function: unknown

Regulator: unknown

Top five genes with highest weights:

| **Locus tag** | **Gene weight** | **Gene name** | **Gene product** |
|---|---|---|---|
| PP_5580 | 0.170614 | *PP_5580* | protein of unknown function |
| PP_5578 | 0.164875 | *PP_5578* | conserved protein of unknown function |
| PP_5579 | 0.154913 | *PP_5579* | conserved protein of unknown function |
| PP_3109 | 0.114233 | *PP_3109* | conserved protein of unknown function |
| PP_5582 | 0.110593 | *PP_5582* | conserved protein of unknown function |

Differentially active condition: *mfsR* overexpression, $H_2O_2$ addition

**70. Citrate iModulon (*n*=5, explained variance: 0.26%)**

Category: Functional iModulon

Inferred function: Citrate transportation

Regulator: unknown

Top five genes with highest weights:

| Locus tag | Gene weight | Gene name | Gene product |
| --- | --- | --- | --- |
| PP_1419 | 0.225185 | *opdH* | tricarboxylate-specific outer membrane porin |
| PP_1417 | 0.213549 | *PP_1417* | putative Tricarboxylate transport protein TctB |
| PP_1418 | 0.200378 | *PP_1418* | putative Tricarboxylate transport protein TctC |
| PP_0147 | 0.130915 | *citN* | Citrate transporter |
| PP_1415 | 0.128599 | *PP_1415* | putative membrane associated enzyme subunit |

Differentially active condition: *mfsR* overexpression, $H_2O_2$ addition

**71. GlcC iModulon (*n*=4, explained variance: 0.25%)**

Category: Regulatory iModulon

Inferred function: Glycolate catabolism

Regulator: GlcC (PP_3744)

Four genes with their weights:

| Locus tag | Gene weight | Gene name | Gene product |
| --- | --- | --- | --- |
| PP_3745 | 0.240653 | *glcD* | glycolate oxidase putative FAD-linked subunit |
| PP_3747 | 0.225169 | *glcF* | glycolate oxidase iron-sulfur subunit |

| | | | |
|---|---|---|---|
| PP_3748 | 0.222878 | *glcG* | conserved hypothetical protein |
| PP_3746 | 0.182284 | *glcE* | glycolate oxidase putative FAD-binding subunit |

Differentially active condition: glycolaldehyde addition, growth phase, *relA* deletion, H2O2 addition

#### 72. VanR iModulon (*n*=4, explained variance: 0.24%)

Category: Regulatory iModulon (MR and RR of 0.75 and 1.00, well-matched)

Inferred function: Aromatic acid catabolism

Regulator: VanR (PP_3738)

Four genes with their weights:

| Locus tag | Gene weight | Gene name | Gene product |
|---|---|---|---|
| PP_3736 | 0.239561 | *vanA* | vanillate O-demethylase oxygenase subunit |
| PP_3737 | 0.229448 | *vanB* | vanillate O-demethylase oxidoreductase |
| PP_3740 | 0.184332 | *PP_3740* | Major facilitator family transporter |
| PP_3739 | 0.142731 | *galP-IV* | porin-like protein |

Differentially active condition: ferulate utilization

#### 73. RbsR iModulon (*n*=5, explained variance: 0.23%)

Category: Regulatory iModulon (MR and RR of 0.71 and 0.71, well-matched)

Inferred function: Ribose, nicotinic acid catabolism

Regulator: RbsR (PP_2457)

Top five genes with highest weights:

| Locus tag | Gene weight | Gene name | Gene product |
|---|---|---|---|
| PP_2457 | 0.138211 | *rbsR* | DNA-binding transcriptional repressor |
| PP_2455 | 0.127392 | *rbsA-I* | ribose ABC transporter - ATP-binding subunit |
| PP_2456 | 0.123227 | *rbsC* | D-ribose ABC transporter - permease subunit |
| PP_1638 | 0.119737 | *fpr-I* | ferredoxin--NADP(+) reductase |
| PP_3944 | 0.11521 | *nicC* | 6-hydroxynicotinate 3-monooxygenase |

Differentially active condition: *finR* deletion, growth phase

### 74. IonicLiquid iModulon (*n*=14, explained variance: 0.21%)

Category: Functional iModulon

Inferred function: Triethylamine hydrogen sulfate stress

Regulator: unknown

Top five genes with highest weights:

| Locus tag | Gene weight | Gene name | Gene product |
|---|---|---|---|
| PP_4930 | 0.205191 | *emrE* | putative small multidrug resistance protein |
| PP_3725 | 0.183095 | *PP_3725* | putative Acyl-CoA dehydrogenase |
| PP_3726 | 0.168649 | *PP_3726* | Enoyl-CoA hydratase/isomerase family protein |
| PP_2817 | 0.161188 | *mexC* | Multidrug efflux RND membrane fusion protein MexC |

| | | | |
|---|---|---|---|
| PP_2816 | 0.14272 | *PP_2816* | putative Transcriptional regulator NfxB |

Differentially active condition: ionic liquid (triethylamine hydrogen sulfate) addition

#### 75. Zinc iModulon (*n*=14, explained variance: 0.21%)

Category: Functional iModulon

Inferred function: Metal (zinc) related response

Regulator: unknown

Top five genes with highest weights:

| **Locus tag** | **Gene weight** | **Gene name** | **Gene product** |
|---|---|---|---|
| PP_0032 | 0.419372 | *PP_0032* | conserved hypothetical protein |
| PP_0046 | 0.237019 | *opdT-I* | tyrosine-specific outer membrane porin D |
| PP_0028 | 0.187564 | *PP_0028* | conserved hypothetical protein |
| PP_0042 | 0.187183 | *PP_0042* | conserved exported protein of unknown function |
| PP_1439 | 0.184691 | *PP_1439* | conserved exported protein of unknown function |

Differentially active condition: zinc ion addition

#### 76. Genomic-3 iModulon (*n*=1, explained variance: 0.17%)

Category: Genomic iModulon

Active condition: HNS study (*turB* deletion)

#### 77. Uncharacterized-4 iModulon (*n*=5, explained variance: 0.16%)

Category: Uncharacterized iModulon

Inferred function: unknown

Regulator: unknown

Top five genes with highest weights:

| Locus tag | Gene weight | Gene name | Gene product |
|---|---|---|---|
| PP_2734 | 0.259891 | *cfa* | Cyclopropane-fatty-acyl-phospholipid synthase |
| PP_2733 | 0.241367 | *PP_2733* | conserved membrane protein of unknown function |
| PP_5173 | -0.18507 | *PP_5173* | RND efflux transporter |
| PP_5174 | -0.19752 | *PP_5174* | Efflux membrane fusion protein%2C RND family |
| PP_5175 | -0.19754 | *PP_5175* | HlyD family secretion protein |

Differentially active condition: 3227 strain specific

**78. Null-3 iModulon (*n*=0)**

**79. PedR1-1 iModulon (*n*=2, explained variance: 0.11%)**

Category: Regulatory iModulon (MR and RR of 1.00 and 0.06, regulon subset)

Inferred function: Alcohol/aromatics catabolism

Regulator: PedR1 (PP_2665)

Two genes with their weights:

| Locus tag | Gene weight | Gene name | Gene product |
|---|---|---|---|
| PP_2674 | 0.371647 | *qedH-I* | quinoprotein ethanol dehydrogenase |
| PP_2673 | 0.257685 | *PP_2673* | Pentapeptide repeat family protein |

Differentially active condition: substrate type, mfsR overexpression

#### 80. Sulfur iModulon (*n*=5, explained variance: 0.16%)

Category: Functional iModulon

Inferred function: element (sulfur) homeostasis

Regulator: unknown

Four genes with their weights:

| **Locus tag** | **Gene weight** | **Gene name** | **Gene product** |
|---|---|---|---|
| PP_3639 | 0.370612 | *PP_3639* | Alkylhydroperoxidase AhpD domain protein |
| PP_3638 | 0.183831 | *PP_3638* | putative Acyl-CoA dehydrogenase |
| PP_3637 | 0.172138 | *PP_3637* | putative Sulfonate ABC transporter%2C ATP-binding protein |
| PP_3636 | 0.162793 | *PP_3636* | putative Sulfonate ABC transporter%2C periplasmic sulfonate-binding protein |

Differentially active condition: *wspR* overexpression

#### 81. Noise-2 iModulon

#### 82. Noise-1 iModulon

#### 83. Noise-4 iModulon

#### 84. Noise-3 iModulon

**Supplementary Note 2. Nucleotide symbols**

| Symbol | Full Name |
|---|---|
| A | Adenine |
| C | Cytosine |
| G | Guanine |
| T | Thymine |
| U | Uracil |
| R | Guanine / Adenine (purine) |
| Y | Cytosine / Thymine (pyrimidine) |
| K | Guanine / Thymine |
| M | Adenine / Cytosine |
| S | Guanine / Cytosine |
| W | Adenine / Thymine |
| B | Guanine / Thymine / Cytosine |
| D | Guanine / Adenine / Thymine |
| H | Adenine / Cytosine / Thymine |
| V | Guanine / Cytosine / Adenine |
| N | Adenine / Guanine / Cytosine / Thymine |

**Supplementary Note 3. The naming of evolved strains**

Strains were previously named according to ALEdb conventions[2], with each separate evolution experiment receiving a different ALE (A) number, which is iteratively assigned new flask (F) numbers as the experiment runs; one isolate (I) was taken and sequenced from each flask of interest.

### Supplementary Methods

#### Supplementary Method 1. Transcriptome sequencing (RNA-seq)

Newly generated RNA-sequencing samples were prepared as reported previously[2,3]. Briefly, cells were cultured in either LB medium (10 g/L tryptone, 5 g/L yeast extract, 10 g/L NaCl) or the modified minimal M9 medium. The minimal medium contains 4 g/L glucose, 2 g/L $(NH_4)_2SO_4$, 6.8 g/L $Na_2HPO_4$, 3 g/L $KH_2PO_4$, 0.5 g/L NaCl, 2 mM $MgSO_4$, 0.1 mM $CaCl_2$, 500 μL/L 2000× trace element solution). The composition of the trace element solution is 4.5 g/L $ZnSO_4 \cdot 7H_2O$, 0.7 g/L $MnCl_2 \cdot 4H_2O$, 0.3 g/L $CoCl_2 \cdot 6H_2O$, 0.2 g/L $CuSO_4 \cdot 2H_2O$, 0.4 g/L $Na_2MoO_4 \cdot 2H_2O$, 4.5 g/L $CaCl_2 \cdot 2H_2O$, 3.0 g/L $FeSO_4 \cdot 7H_2O$, 1.0 g/L $H_3BO_3$, 0.1 g/L KI, 15 g/L disodium ethylenediaminetetraacetate. Colonies grown on LB agar plates were inoculated into 15 mL of the M9 medium and incubated at 30 °C with continuous stirring at 1,100 rpm. After overnight, cultures were diluted into the fresh medium at $OD_{600}$ of 0.05. When $OD_{600}$ reached ~1, the cultures were further diluted into a new fresh medium at $OD_{600}$ of 0.05. Once $OD_{600}$ reached 0.6-0.8, 3 mL of cultures were directly mixed with 6 mL of an RNAprotect Bacteria Reagent from Qiagen (Hilden, Germany). Subsequently, the total RNA was extracted by using a Quick-RNA Fungal/Bacterial Miniprep kit from Zymo Research (Irvine, CA, USA). Ribosomal RNA was removed from total RNA samples using anti-ribosomal oligonucleotides as described[3]. Sequencing libraries were prepared by using a KAPA RNA HyperPrep kit from Kapa Biosystems (Wilmington, MA, USA) or a SWIFT Rapid RNA Library Kit from Swift Biosciences (Ann Arbor, MI, USA) by following manufacturers' protocols. Libraries were sequenced using an illumina NovaSeq 6000 at the UC San Diego IGM Genomics Center.

The 12 samples for the "aromatic" project were prepared with several modifications. Cells were grown in the M9 medium with 2.5 g/L of either coumarate, ferulate, a mixture of coumarate and ferulate, or glucose. Collected cells were resuspended in TRIzol$^{TM}$ reagent (ThermoFisher Scientific, Waltham, MA, USA). Total RNA was isolated using the RNeasy Mini Kit (Qiagen) by following the manufacturer's instructions. Ribosomal RNA was removed from total RNA using the Ribo-Zero rRNA removal kit (illumina). Sequencing libraries were prepared using the ScriptSeq$^{TM}$ Complete kit (illumina). Library construction and sequencing were conducted by the Genomics Core Facility at the University of Nebraska Medical Center.

**Supplementary Method 2. Determination of iModulon threshold values**

For a given iModulon, its member genes were defined by screening a threshold value (i.e., cut-off) from the **M** matrix. In detail, the **M** matrix contains weights of every gene for an iModulon where most genes have nearly zero values and a small number of meaningful genes have high weights. An optimal threshold value was determined by performing the D'Agostino $K^2$ test of scikit-learn[4]. For each iModulon, the gene with the highest absolute gene weight was iteratively removed and the $\mathrm{K}^2$ value was calculated. When non-removed genes sufficiently displayed the normal distribution around zero, the value was chosen as a threshold. For the PplR1, Sulfur, FleQ/AmrZ, PP_2260-3, GcsR, PP_0615-8 iModulons, their threshold values were manually adjusted to 0.08, 0.132, 0.053, 0.140, 0.11, 0.11, respectively, to include genes in the same regulons or TUs.

### Supplementary Tables

### Supplementary Table 1. Selected 30 Regulatory iModulons[a]

| iModulon name | Involved regulators | Reported inducers (effectors)[b] | Type | Roles | iM Size | Exp Var | Ref |
|---|---|---|---|---|---|---|---|
| PydR/RpoS | PydR, RpoS | N/A | Regulon discovery | Pyrimidine degradation | 30 | 5.4 | [5] |
| TurA | TurA, TurB | N/A | Regulon discovery | Biosynthesis of lipids nonapeptide phytotoxins | 17 | 4.0 | [6] |
| BkdR | BkdR | Branched amino acids | Regulon discovery | Starvation response | 19 | 2.8 | [7,8] |
| PedR1-2 | PedR1 | Chloramphenicol | Well-matched | Alcohol/aromatics catabolism | 31 | 2.6 | [9,10] |
| Zur | Zur | Zinc ion | Regulon discovery | Metal homeostasis | 33 | 1.9 | [11] |
| PcaR | PcaR, PP_3359, VanR | Aromatic compounds | Regulon discovery | Aromatic acid catabolism | 45 | 1.6 | [12,13] |
| FleQ/AmrZ | FleQ, AmrZ | c-di-GMP | Regulon subset | Flagella synthesis | 44 | 1.3 | [14,15] |
| FliA | FliA, FleQ | N/A | Regulon discovery | Cell mobility | 29 | 1.2 | [14,16] |
| PtxS | PtxS, GnuR | 2-KG, gluconate, 6-PG | Well-matched | Sugar catabolism | 11 | 1.2 | [17,18] |
| PplR1 | PplR1, FleQ | Light | Well-matched | Light inducible iModulon | 22 | 0.9 | [19] |
| GltR-II | GltR-II, Crc | 6-PG, 2-KG | Well-matched | Sugar catabolism | 7 | 0.9 | [18,20] |
| LiuR | LiuR, RbsR, BkdR | Branched amino acids, leucine | Regulon subset | Ribose and lipid catabolism | 12 | 0.8 | [8] |
| FleQ | FleQ, Fur | c-di-GMP | Regulon subset | Inorganic ion transport and metabolism | 36 | 0.8 | [15] |
| GlnG | GlnG | Nitrogen | Regulon subset | Amino acid transportation | 11 | 0.7 | [21] |
| GclR | GclR, RutR-I, GlpR | Allantoin | Regulon subset | C2 carbon metabolism | 14 | 0.7 | [22] |
| GcsR | GcsR | Glycine | Well-matched | Glycine metabolism | 4 | 0.7 | [23] |
| OxyR | OxyR | Reactive oxygen | Regulon discovery | Oxidative stress | 12 | 0.7 | [24–26] |
| HexR | HexR | KDPG | Well-matched | Sugar catabolism | 9 | 0.6 | [18] |
| FruR | FruR | F-1-P | Regulon | Fructose catabolism | 4 | 0.5 | [27] |

| | | | | | | | |
|---|---|---|---|---|---|---|---|
| | | | subset | | | | |
| AmrZ | AmrZ, FleQ | c-di-GMP | Well-matched | Alginate biosynthesis | 12 | 0.5 | 28,29 |
| BenR | BenR, CatR | Muconate | Well-matched | Aromatic acid catabolism | 10 | 0.5 | 30,31 |
| AcoR | AcoR | N/A | Well-matched | Acetoin catabolism | 5 | 0.5 | N/A |
| HutC | HutC | cis-Urocanic acid | Well-matched | Histidine metabolism | 9 | 0.4 | 32 |
| PP_5350 | PP_5350 | Acetate | Regulon subset | Glyoxylate and dicarboxylate metabolism | 4 | 0.4 | 3 |
| Metr-I | Metr-I | Homocysteine | Regulon subset | Homocysteine metabolism | 6 | 0.4 | 33 |
| LldR | LldR, Crc | L-lactate | Well-matched | Lactate catabolism | 4 | 0.3 | 34 |
| GlcC | GlcC | Glycolate | Well-matched | Glycolate catabolism | 4 | 0.2 | 35 |
| VanR | VanR | Vanillate | Well-matched | Aromatic acid catabolism | 4 | 0.2 | 36 |
| RbsR | RbsR, FinR | Ribose | Regulon subset | Ribose, nicotinic acid catabolism | 7 | 0.2 | N/A |
| PedR1-1 | PedR1 | Chloramphenicol | Regulon subset | Alcohol/aromatics catabolism | 2 | 0.1 | 9,10 |

[a]Well-matched, regulon subset, regulon discovery iModulons were listed in decreased order of their explained variances.

[b]Adapted from RegPrecise (http://regprecise.sbpdiscovery.org:8080/WebRegPrecise/index.jsp) or listed related references.

Abbreviations: c-di-GMP, cyclic diguanylate; 2-KG, 2-ketogluconate; 6-PG, 6-phosphogluconate; F-1-P, Fructose 1-phosphate; KDPG, 2-dehydro-3-deoxy-phosphogluconate; N/A, not available.

**Supplementary Table 2. The best enriched motif from transcriptional units in iModulons**

| Name[a] | Enriched motif[b] | E-value[c] | # of TUs[d] | # of genes[e] | Width |
|---|---|---|---|---|---|
| ColR | 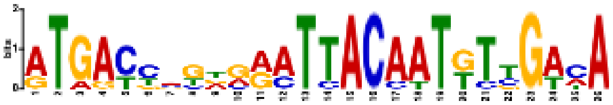 ATGACYMKDRRATTACAATKTYGAMA | 1.1 $\times 10^{-11}$ | 7/8 88% | 9/11 82% | 26 |
| Exporters | 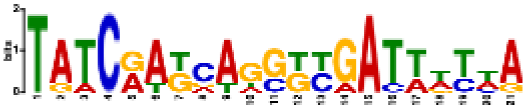 TATCRAKCAGSKYGATWWYWA | 5.3 $\times 10^{-4}$ | 6/8 75% | 8/10 80% | 21 |
| FleQ/ AmrZ | 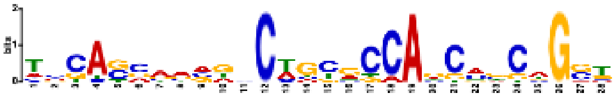 TMCASSAAVDNCWGYRCCAGCMBCWGST | 1.5 $\times 10^{-7}$ | 18/18 100% | 44/44 100% | 28 |
| FleQ/ Fur* | 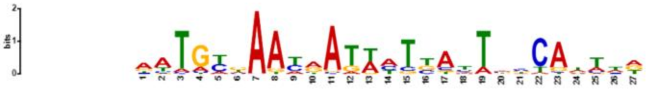 AATGYKAAYRATWHTBAYTHBCAWTWD 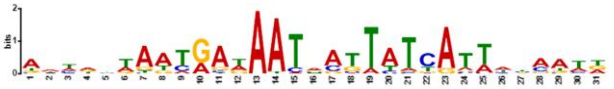 Fur in *E. coli* (E-value: 2.51$\times 10^{-7}$, database: collectTF) | 1.7 $\times 10^{-46}$ | 25/27 93% | 34/36 94% | 27 |
| FliA | 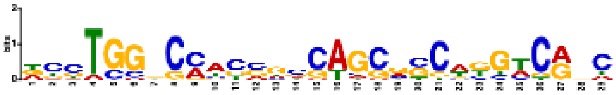 DYCTGGVCMWCSGCCAGCRSCWKGWCRNC | 7.2 $\times 10^{-7}$ | 21/25 84% | 24/29 83% | 29 |
| FnrA-1 | 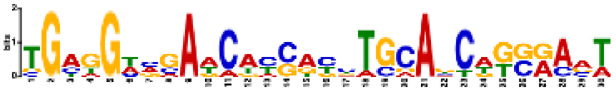 TGAGGWVGAWCAYSAYYTGCAKCWKSRAVT | 3.0 $\times 10^{-6}$ | 8/10 80% | 14/16 88% | 30 |
| FnrA-2* | 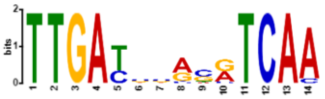 TTGAYSSRSRTCAA 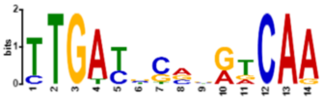 Anr in *P. aeruginosa* (E-value: 3.39$\times 10^{-8}$, database: prodoric) | 3.9 $\times 10^{-17}$ | 12/12 100% | 18/18 100% | 14 |

| | | | | | |
|---|---|---|---|---|---|
| GbdR | 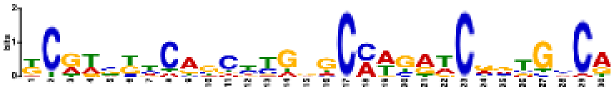 KCRWBKYCAGCYYGBGCMWGRTCRVTGSCA | 2.4 $\times10^{-17}$ | 20/24 83% | 28/36 78% | 30 |
| GclR | 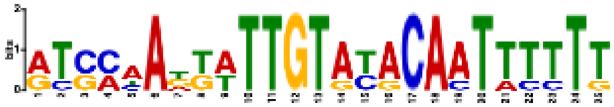 RTSMAADKWTTGTAYACAATTTTTT | 4.2 $\times10^{-9}$ | 5/10 50% | 8/14 57% | 29 |
| HexR* | 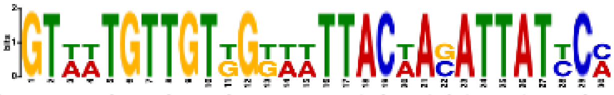 GTWWTGTTGTKGKWWTTACWASATTATYCM | 7.5 $\times10^{-15}$ | 4/4 100% | 9/9 100% | 30 |
| Ionic Liquid | 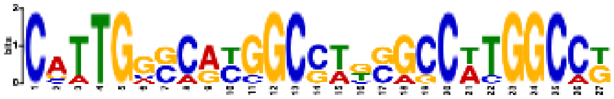 CATTGGSCRYGGCSWKSGCCWTGGCCK | 4.3 $\times10^{-8}$ | 6/9 67% | 7/14 50% | 27 |
| LiuR | 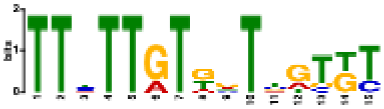 TTYTTGTKRTYGTKT | 8.0 $\times10^{-5}$ | 8/8 100% | 12/12 100% | 15 |
| Multiple stress-1* | 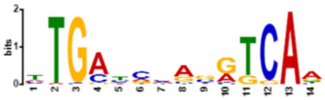 TTGAYCHASRTCAA 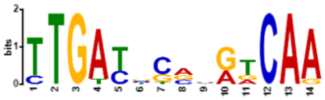 Anr in *P. aeruginosa* (E-value: $1.89\times10^{-6}$, database: prodoric) | 5.1 $\times10^{-23}$ | 48/48 100% | 58/58 100% | 14 |
| Multiple stress-2 | 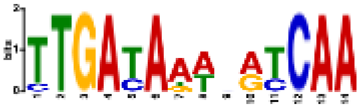 TTGATAAWNRTCAA | 2.3 $\times10^{-7}$ | 7/9 78% | 7/9 78% | 14 |
| Osmotic stress-1 | 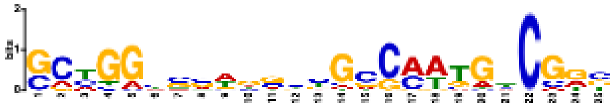 SCYGGYSVWGSHBGCCMWTGBCGRB | 9.8 $\times10^{-12}$ | 39/43 91% | 42/46 91% | 25 |
| Osmotic stress-2 | 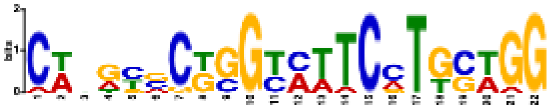 CWNRYVCKGGYMTTCMTGSWGG | 1.7 $\times10^{-8}$ | 10/23 43% | 12/31 39% | 22 |

| | | | | | |
|---|---|---|---|---|---|
| PcaR | 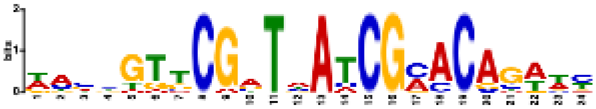<br>DASHGTTCGATWATCGMACAGWWY | $7.0\times10^{-20}$ | 15/32<br>47% | 21/45<br>47% | 24 |
| PedR1-2 | 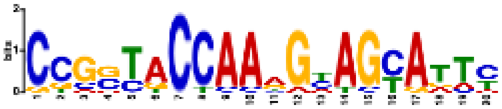<br>CCGSTACCAARGYAGYATTY | $3.0\times10^{-11}$ | 11/17<br>65% | 20/31<br>65% | 20 |
| Phage-2 | 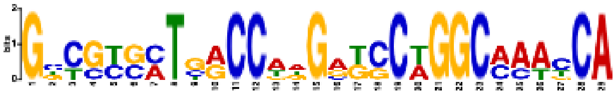<br>GBCGYSMTSRCCWDGRKCCWGGCMAAYCA | $1.2\times10^{-11}$ | 7/20<br>35% | 12/47<br>26% | 29 |
| PP_0204<br>/CysB | 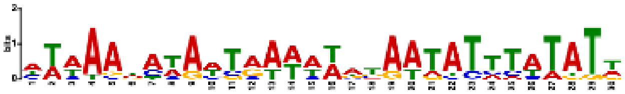<br>HWWAAHAWAWTRAWWWATAATATTTATAT<br>W | $3.5\times10^{-21}$ | 11/17<br>65% | 23/29<br>79% | 29 |
| PplR1 | 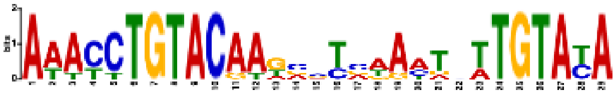<br>AAACCTGTACAAKVSTVWAMTNTTGTAYA | $2.0\times10^{-26}$ | 7/9<br>78% | 19/22<br>86% | 29 |
| Putrescine | 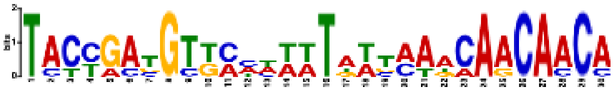<br>TACYGATGTKMHHWWTAWTMAACAACAACA | $5.7\times10^{-6}$ | 5/6<br>83% | 6/7<br>86% | 30 |
| PvdS* | 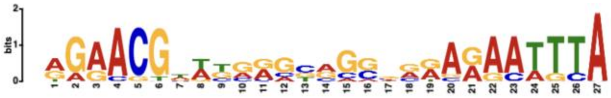<br>WWAYRAHAWTKAATATCAKYHT<br>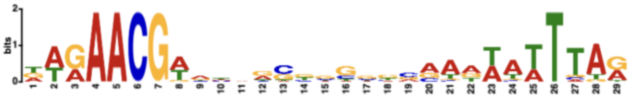<br>PvdS in *P. syringae*<br>(E-value: $2.19\times10^{-14}$, prodoric) | $6.7\times10^{-12}$ | 11/19<br>58% | 16/36<br>44% | 27 |
| PydR/<br>RpoS* | 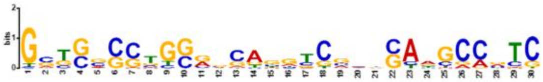<br>GSTSSSSTGSRVCAGGTCSVNSAAGCCVTC<br>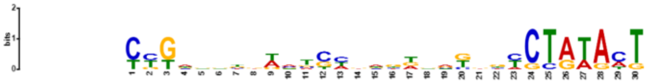<br>RpoS in *E. coli*<br>(E-value: $8.58\times10^{-4}$, database: dpinteract) | $2.8\times10^{-7}$ | 27/27<br>100% | 30/30<br>100% | 30 |
| TCA<br>cycle* | 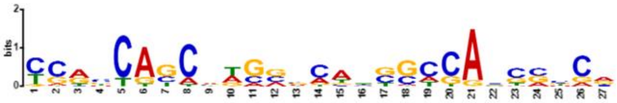 | $1.8\times10^{-11}$ | 49/50<br>98% | 54/55<br>98% | 27 |

| | | | | | |
|---|---|---|---|---|---|
| | YCRSCAGCVWGVVCABSSCCABCSCCR 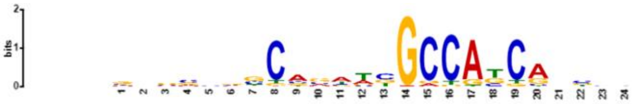 AmrZ in *P. aeruginosa* (E-value: 2.79×$10^{-5}$, database: collectTF) | | | | |
| Translation | 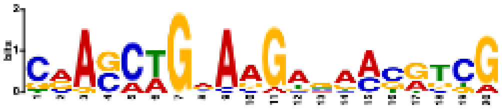 CRASCWGVAAGAGAMSRTCG | 2.6 ×$10^{-4}$ | 14/16 88% | 30/45 67% | 20 |
| TurA-1 | 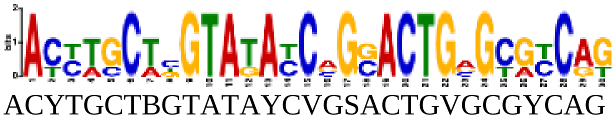 ACYTGCTBGTATAYCVGSACTGVGCGYCAG | 8.2 ×$10^{-4}$ | 4/6 67% | 12/17 71% | 30 |
| Unchar-5 | 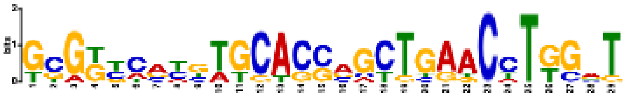 GCGKYCVYBTGCACSVGCTGAACCTKGRT | 1.4 ×$10^{-5}$ | 8/12 67% | 8/12 67% | 29 |
| Unchar-7 | 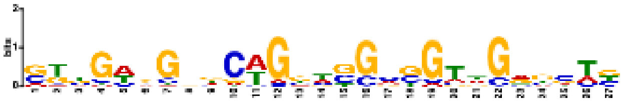 SKBGWBGNDCWGGWSGMSGTWGRKSTB | 2.1 ×$10^{-5}$ | 41/67 61% | 73/117 62% | 27 |
| Unchar-8 | 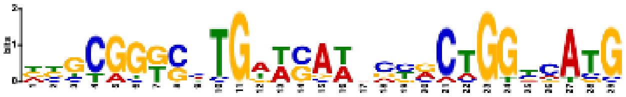 DBGCGGKSMTGWWSAWNSYRCTGGYCATG | 1.9 ×$10^{-16}$ | 11/11 100% | 14/14 100% | 29 |
| Zinc | 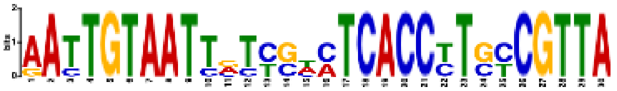 AATTGTAATTVTYSHMTCACCYTSYCGTTA | 5.0 ×$10^{-17}$ | 5/15 33% | 5/17 29% | 30 |
| Zur* | 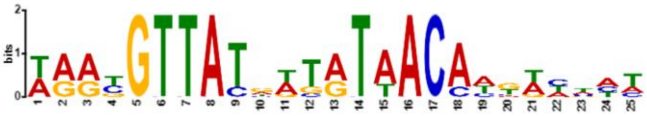 WRRYGTTATRWTRTAACAWKWHWHW 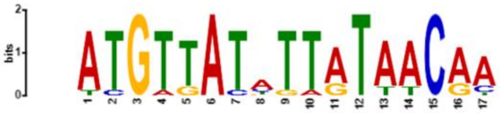 Zur in *P . protegens* (E-value: 1.39×$10^{-5}$, database: collectTF) | 1.4 ×$10^{-27}$ | 11/19 58% | 22/33 67% | 25 |

[a]iModulons are listed alphabetically. Asterisk(*) indicates that an enriched motif matches a previously reported motif in a public database using TOMTOM[37].

[b]If there is a matched motif, both enriched and reported motifs were shown. An E-value

indicating their similarity and the source of a reported motif were additionally given.

[c]The statistical significance of an identified motif by using MEME[38].

[d]The number of transcription units (TUs) which the enriched motif was found from. The number after a slash (/) indicates the total number of TUs in an iModulon.

**Supplementary Table 3. iModulons with differential activity changes between the exponential growth phase and the stationary phase**

| iModulon name | Activity change | Note |
|---|---|---|
| ***Expected changes (consistent to literature or apparent)*** | | |
| PydR/RpoS | 51.9 | Containing genes for pyrimidine degradation and stress response. Its increased activity is due to nucleotide breakdown and reassimilation[5]. |
| LiuR | 23.8 | Containing genes for ribose and lipid catabolism. Its increased activity is due to the depletion of carbon sources and reassimilation of cellular components. |
| AA transport | 14.6 | Containing genes for (active) transport of amino acids. L- or D- Lysine has been suggested as an inducer[39]. Its increased activity is due to the depletion of the major carbon source and a necessity for the reassimilation of amino acids. |
| FnrA-1 | 14.3 | Containing genes known to be regulated by an oxygen-dependent regulator FnrA. Some genes encode oxidases with high affinities to oxygen[40]. Increased activity was due to oxygen limitation at the stationary phase. |
| RbsR | 13.6 | Containing genes for ribose and nicotinic acid catabolism. Increased activity is due to the depletion of the major carbon source. |
| GcsR | 10.8 | Containing genes for glycine or serine metabolism. Increased activity is due to the depletion of the major carbon source. |
| PP_0613-20 | 8.3 | Containing genes for branched amino acid transportation. Increased activity is due to the depletion of the major carbon source. |
| PP_2034-7 | 8.2 | Containing genes for benzoate catabolism. Increased activity is likely due to the depletion of the major carbon source. |
| Putrescine | 6.2 | Containing genes for putrescine metabolism. Putrescine metabolism genes are known to be up-regulated under stressed conditions with a close association to RpoS[41]. |
| OxyR | 5.0 | Containing genes for mitigating oxidative stress. Similar genes are known to be up-regulated by the activity of RpoS[42]. |
| LldR | -6.4 | Containing genes for lactate utilization/production. Decreased activity is likely due to reduced glucose consumption. |
| FruR | -7.0 | Containing genes for fructose catabolism[27]. |

| | | |
|---|---|---|
| HexR | -8.2 | Containing genes for hexose utilization[18]. Decreased activity is likely due to reduced glucose consumption |
| InfA | -17.4 | Containing genes for protein synthesis. |
| Translation | -39.4 | Containing genes for protein synthesis. |
| ***Unexpected changes*** | | |
| BkdR | 28.6 | Containing genes for a starvation response, branched amino acid metabolism[7,8]. However, the functions of most high-weight genes are unknown. |
| Osmotic stress-1 | 11.6 | Containing genes for osmotic stress response genes encoding lipoproteins, membrane components. Although increased osmotic stress response at the stationary phase was observed in *E. coli*[43], their relationship is still not clear. In addition, this iModulon contains a lot of genes with unknown functions. |
| GbdR | 10.9 | Containing genes for catabolism of glycine betaine, osmoprotectant. Its high activity is likely due to increased osmotic stress response. |
| FliA | 7.5 | Containing genes for flagella synthesis and chemotaxis. Its increased activity was unexpected since flagella synthesis is known to be down-regulated during starvation conditions in other microorganisms to save cellular resources[44]. |
| TurA-2 | -16.3 | Containing chaperone genes required for protein refolding. Typically, chaperon genes are up-regulated in stressed conditions but its activity was decreased in *P. putida*. |

The two uncharacterized iModulons (Uncharacterized-3 and Uncharacterized-9) were omitted in this Table.

### Supplementary Figures

**Supplementary Figure 1. General information of collected RNA-seq samples**

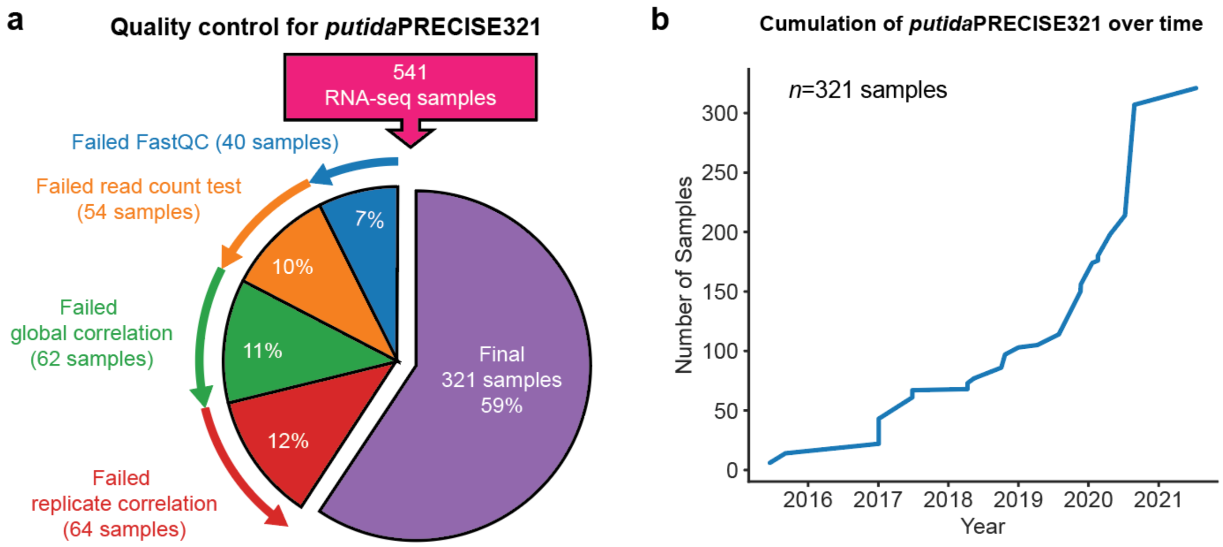


(**a**) Quality control process to generate *putida*PRECISE321 from 541 *P. putida* RNA-seq samples. 40 Samples were discarded owing to failure in their processing with the pipeline or passing FastQC; 54 samples were discarded because of too low read counts; 62 samples were discarded for consistency because they showed poor global correlation or were prepared with non-KT2440 strains; 64 additional samples with poor replicate correlation (<0.95) were discarded. Finally, 321 RNA-seq samples that passed the quality control were utilized as *putida*PRECISE321. (**b**) The cumulative number of the 321 *putida*PRECISE321 samples over time.

**Supplementary Figure 2. Principal component analysis of *putida*PRECISE321**

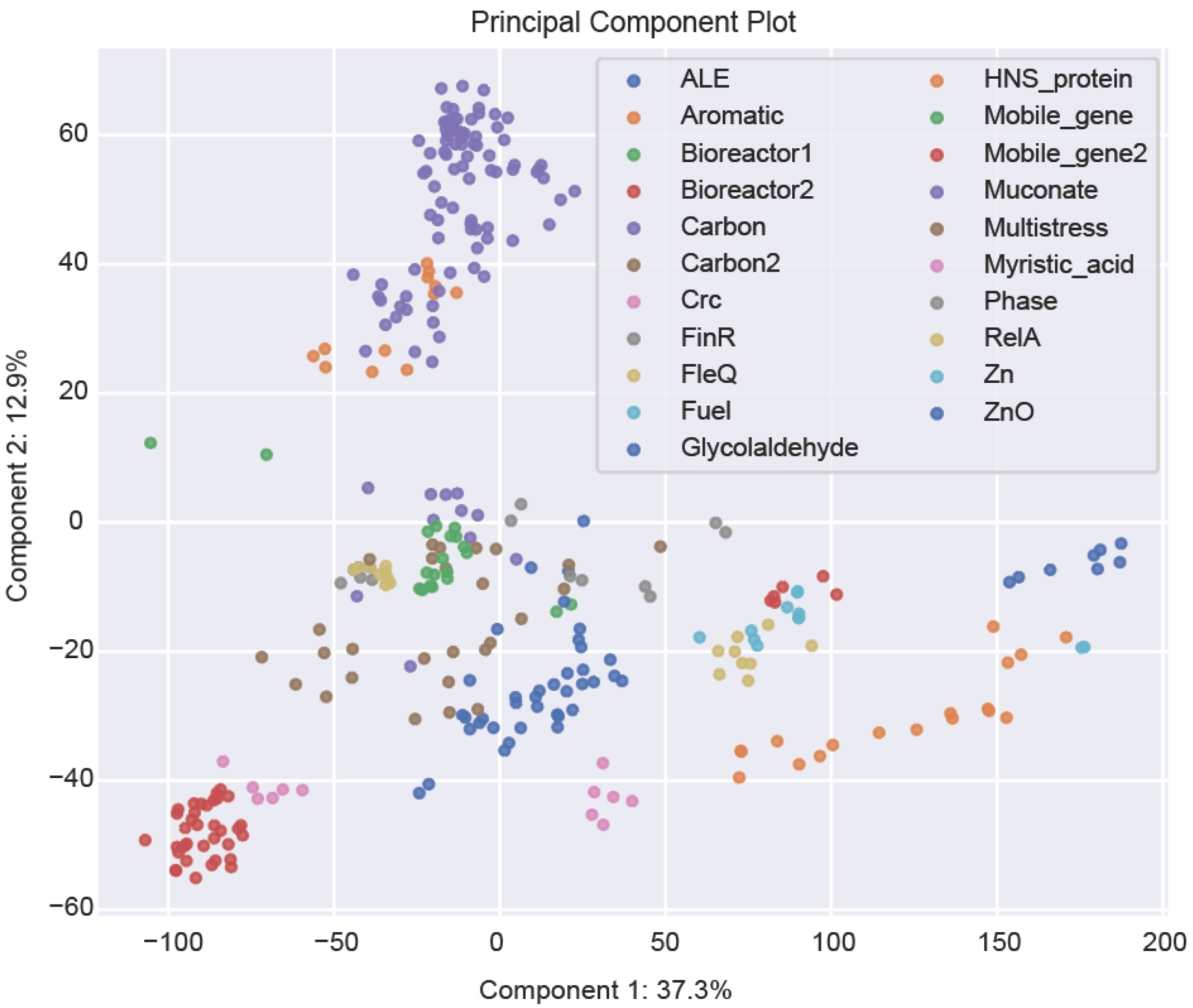


Principal component analysis of *putida*PRECISE321 containing gene expression profiles from 21 projects under 118 conditions.

**Supplementary Figure 3. Characteristics of the 84 *P. putida* iModulons**

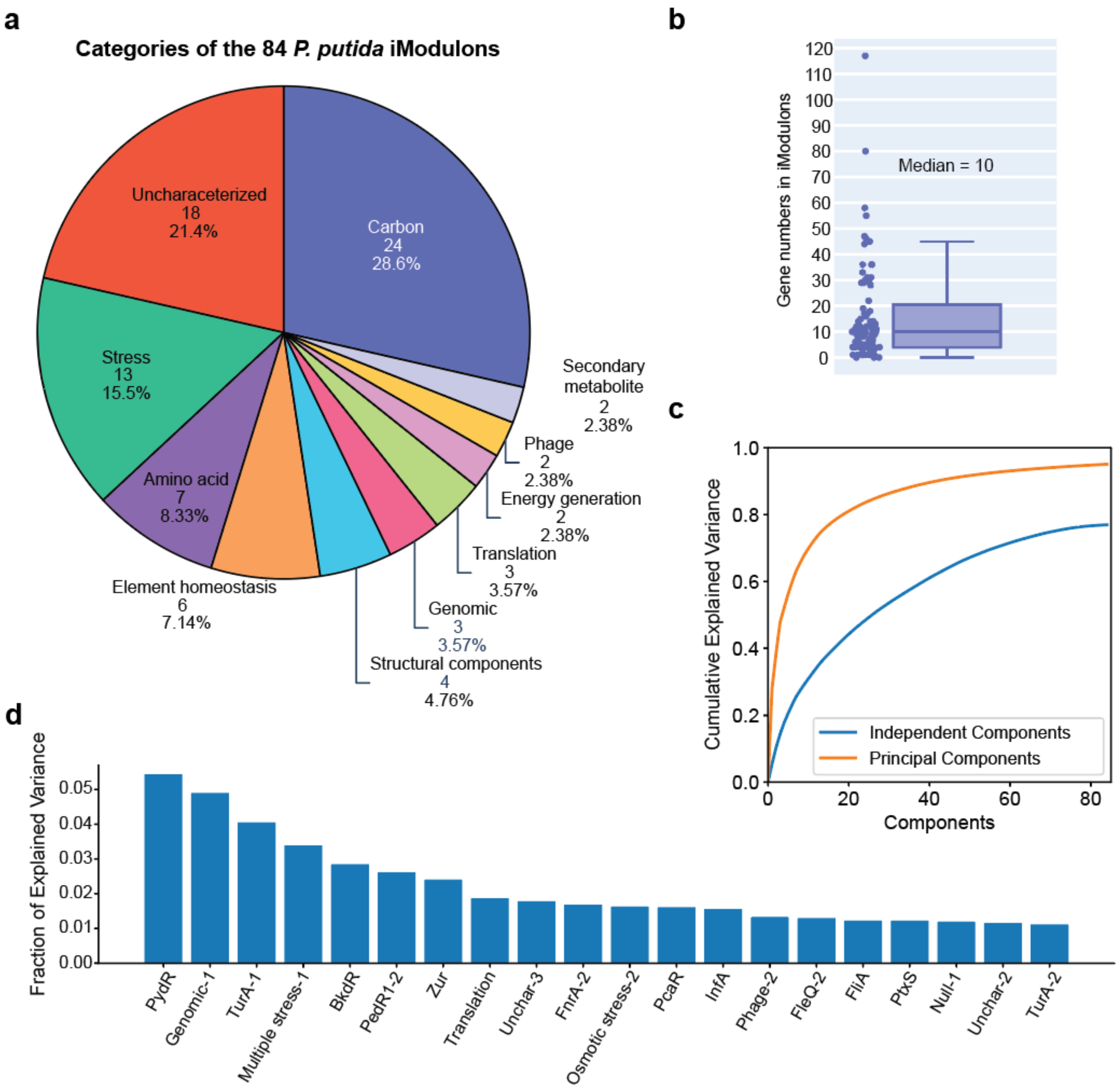


(**a**) Gene number of each iModulon (median: 10) (**b**) Numbers and functions of iModulons (**c**) Cumulative explained variance by independent components (i.e., iModulons) and principal components. (**d**) Fraction of explained variance by top 20 iModulons.

**Supplementary Figure 4. Types of 39 regulatory iModulons depending on iModulon and Regulon recalls**

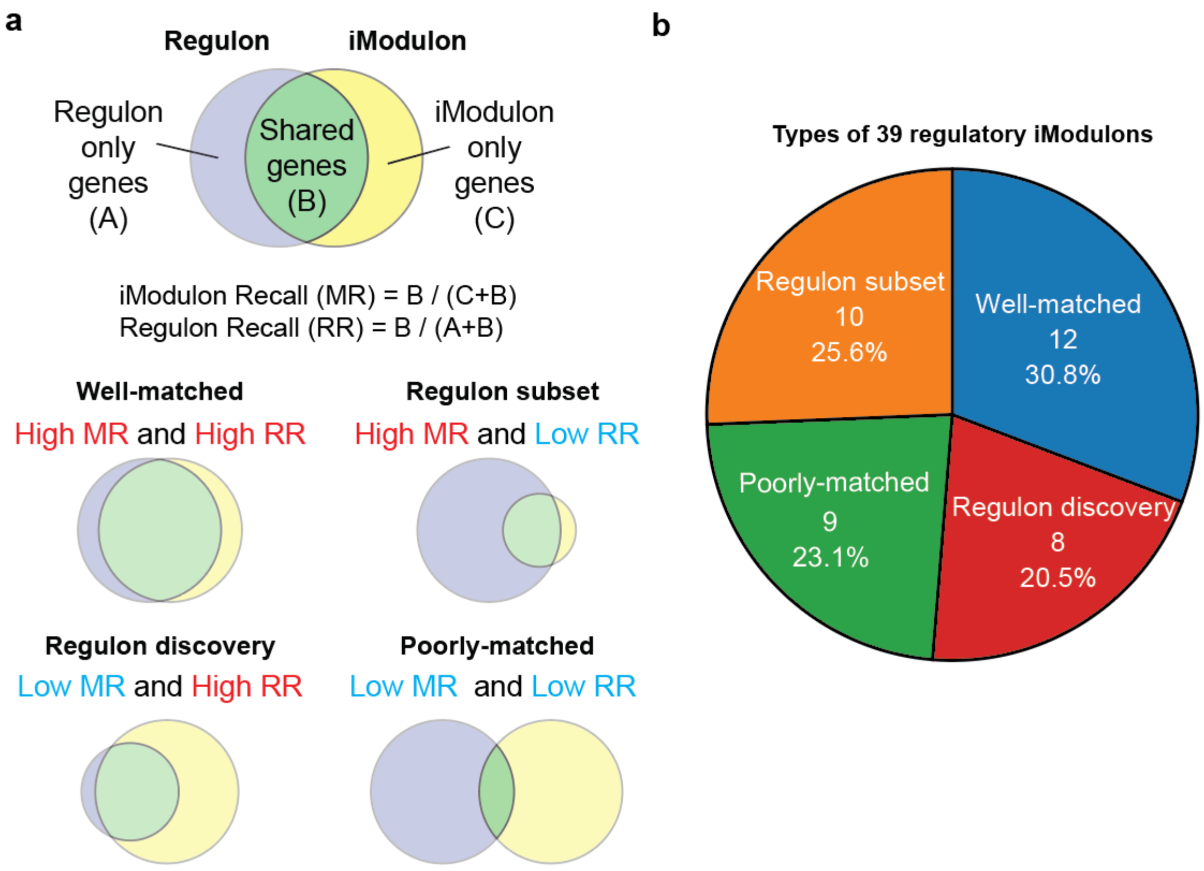


(a) iModulon Recall (MR) and Regulon Recall (RR) were renamed from recall and precision[45]. (b) 4 Types of 39 regulatory iModulons: well-matched (both high MR and RR), regulon subset (high MR but low RR), regulon discovery (low MR and high RR), poorly-matched (both low MR and RR).

**Supplementary Figure 5. Characteristics of the two FleQ-related iModulons (FleQ/Fur and FleQ/AmrZ)**

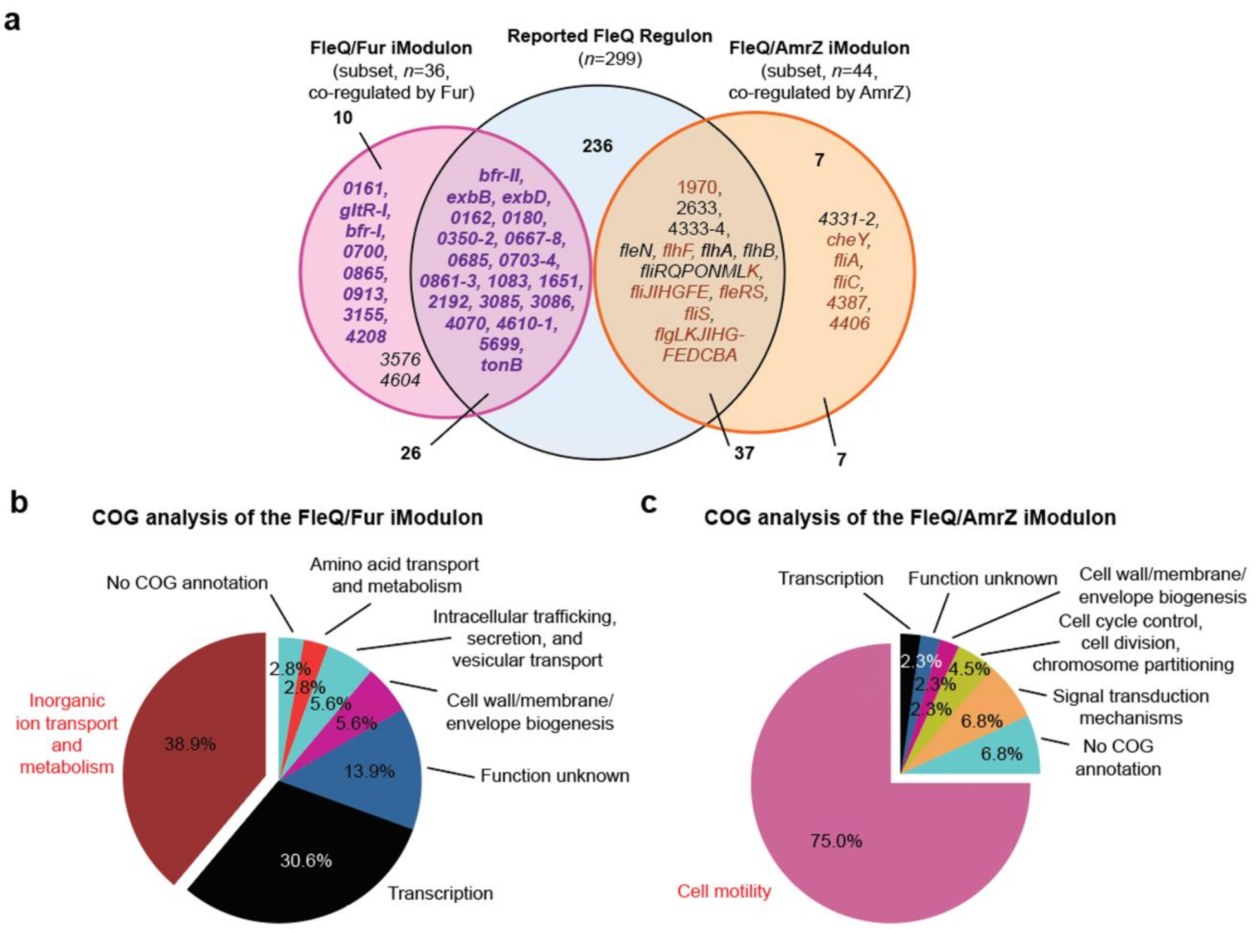


(**a**) A Venn diagram of the FleQ/Fur and FleQ/AmrZ iModulon and the FleQ regulon (**b and c**) Clusters of Orthologous Groups (COG) analysis of the (**b**) FleQ/Fur and (**c**) FleQ/AmrZ iModulons.

**Supplementary Figure 6. Comparison of CysB from *E. coli* K-12 MG1655 and *P. putida* KT2440**

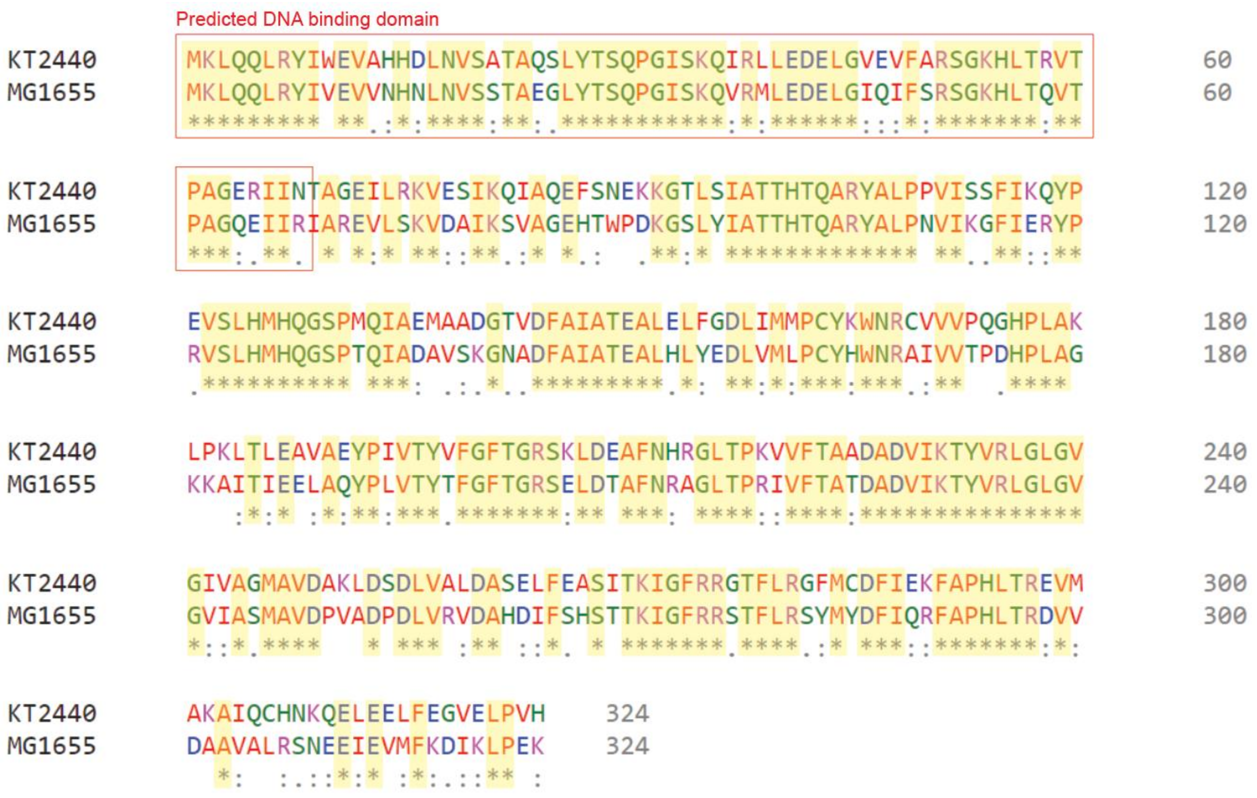


Comparison of the CysB amino acid sequences in *E. coli* K-12 MG1655 (top) and *P. putida* KT2440 (bottom) using CLUSTAL O(1.2.4)[46]. The two sequences share an identity of 63.89%. Red square indicates a predicted DNA binding domain.

**Supplementary Figure 7. Differentially expressed gene analysis of the transcriptomes at the exponential and stationary phase**

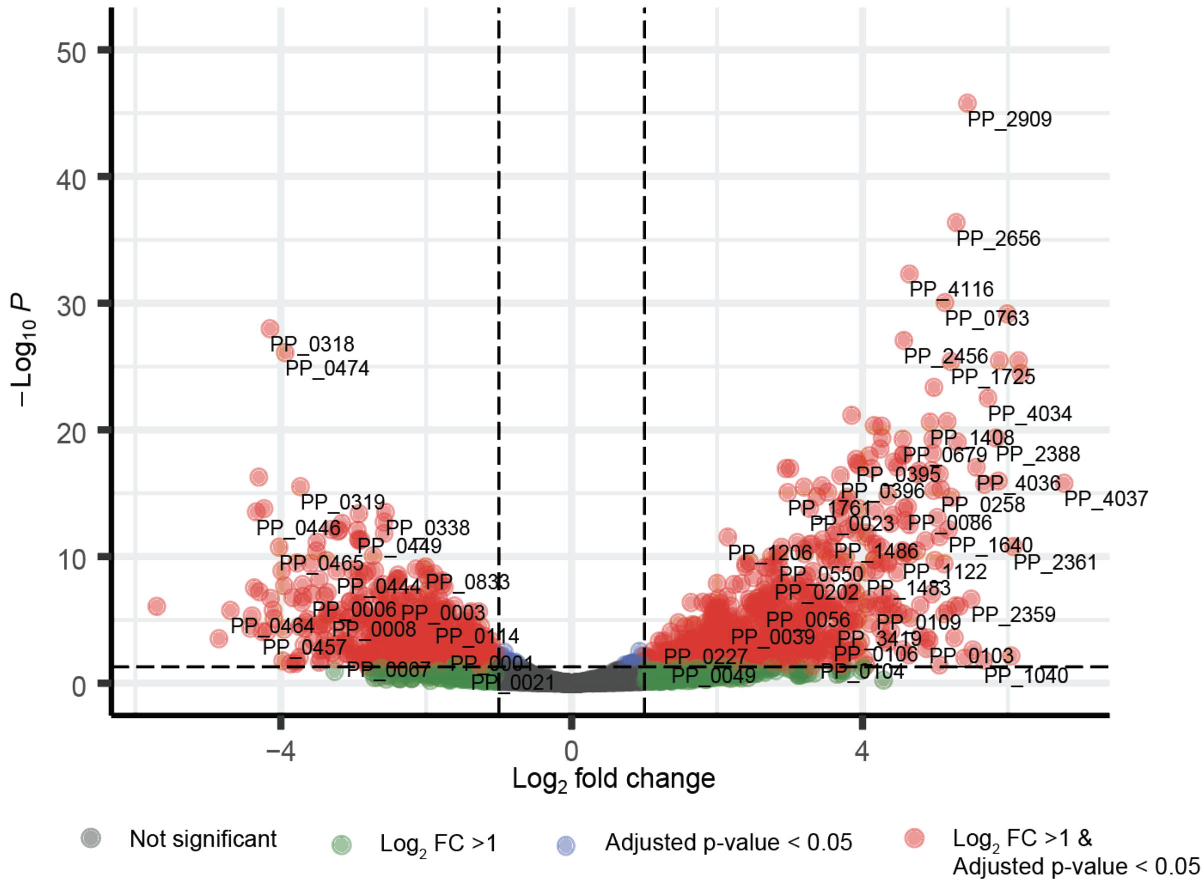


A volcano plot for gene expression changes in the exponential growth and stationary phases (project:*Phase*). Symbols: grey, genes with not significant changes; green, genes with changed expression greater than 2-fold; blue, genes with significantly changed expression ($p_{adj} < 0.05$) but less than 2-fold; red, differentially expressed genes (DEGs) with significantly changed expression ($p_{adj} < 0.05$) and greater than 2-fold. A total of 1454 DEGs were obtained.

**Supplementary Figure 8. Gene weights of the PydR/RpoS and Translation iModulons**

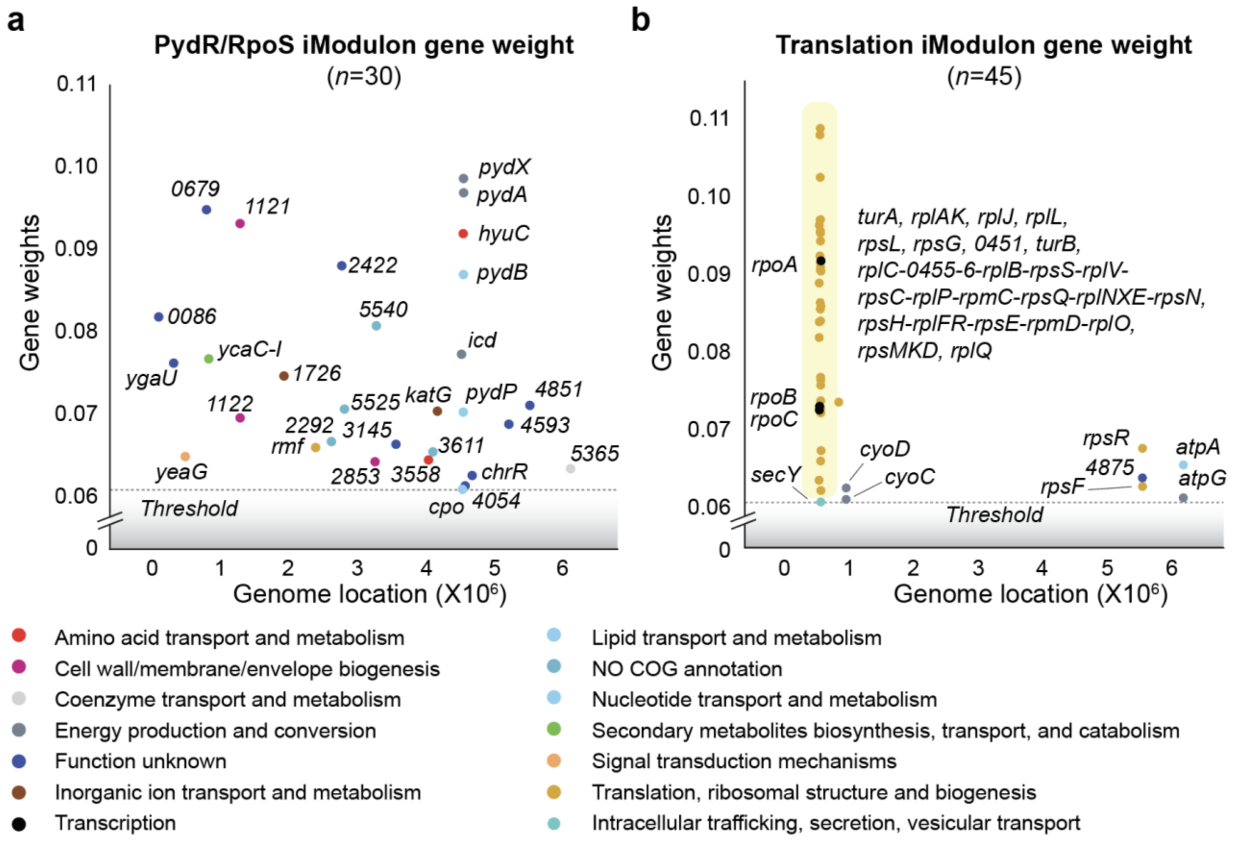


Gene weights of the (**a**) PydR/RpoS and (**b**) Translation iModulons. Colors indicate COG categories. Each scatter plot shows relative gene expression changes compared to the respective reference conditions.

**Supplementary Figure 9. Correlation between the activity of PydR/RpoS iModulon and the expression level of either *pydR* and *rpoS***

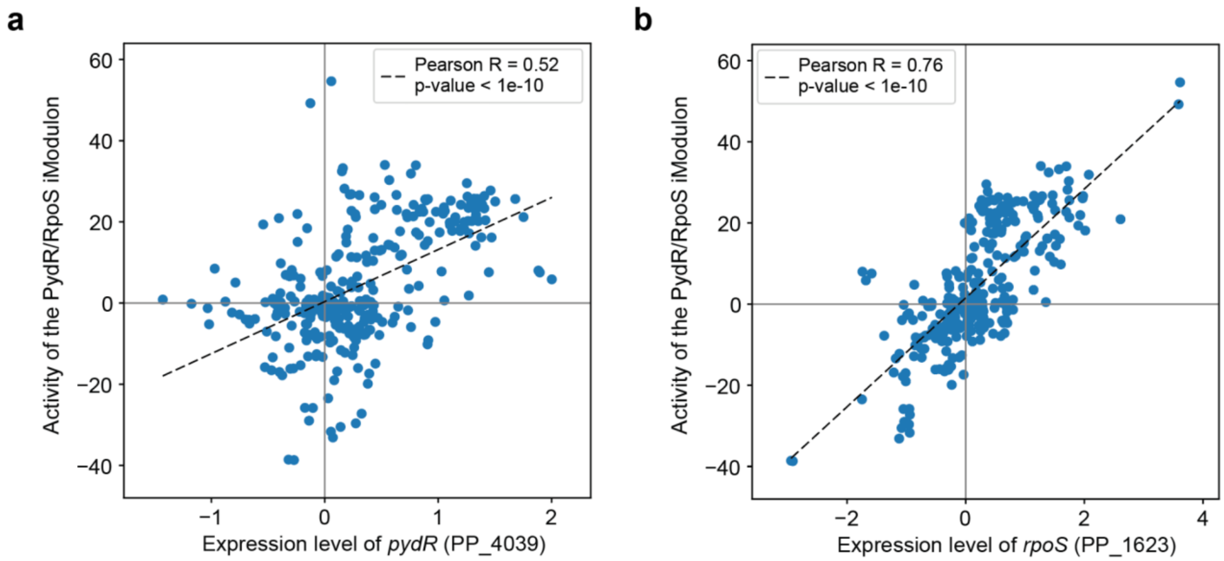


Scatter plots of correlation of the activity of the PydR/RpoS iModulon and the expression of either (**a**) *pydR* (PP_4039) or (**b**) *rpoS* (PP_1623). Dashed lines are the best fits of each plot.
